# Linear integration of sensory evidence over space and time underlies face categorization

**DOI:** 10.1101/2020.11.27.396705

**Authors:** Gouki Okazawa, Long Sha, Roozbeh Kiani

## Abstract

Visual object recognition relies on elaborate sensory processes that transform retinal inputs to object representations, but it also requires decision-making processes that read out object representations and function over prolonged time scales. The computational properties of these decision-making processes remain underexplored for object recognition. Here, we study these computations by developing a stochastic multi-feature face categorization task. Using quantitative models and tight control of spatiotemporal visual information, we demonstrate that humans categorize faces through an integration process that first linearly adds the evidence conferred by task-relevant features over space to create aggregated momentary evidence, and then linearly integrates it over time with minimum information loss. Discrimination of stimuli along different category boundaries (e.g., identity or expression of a face) is implemented by adjusting feature weights of spatial integration. This linear but flexible integration process over *space* and *time* bridges past studies on simple perceptual decisions to complex object recognition behavior.

## Introduction

Accurate and fast discrimination of visual objects is essential to guide our behavior in complex and dynamic environments. Previous studies largely focused on the elaborate sensory mechanisms that transform visual inputs to object-selective neural responses in the inferior temporal cortex of the primate brain through a set of representational changes along the ventral visual pathway (DiCarlo and Cox 2007; Freiwald and Tsao 2010; Riesenhuber and Poggio 1999; Yamins et al. 2014). However, goal-directed behavior also requires decision-making processes that can flexibly read out sensory representations and guide actions based on them as well as information about the environment, behavioral goals, and expected costs and gains. Such processes have been extensively examined using simplified sensory stimuli that vary along a single dimension, e.g., direction of moving dots changing from left to right (Palmer et al. 2005). For those stimuli, subjects’ behavior could be successfully accounted for by flexible mechanisms that accumulate sensory evidence and combine it with task-relevant information (Gold and Shadlen 2007; Ratcliff and Rouder 1998). However, more complex visual decisions based on stimuli defined by multiple features, such as object images, remain underexplored, although the need for such tests is gaining significance and important steps are taken in this direction (Heekeren et al. 2004; Philiastides et al. 2014; Philiastides and Sajda 2006; Zhan et al. 2019).

Here, we apply the quantitative approach developed for studying simple perceptual decisions to investigate face recognition. We focus on face recognition because it is by far the most extensively studied among the sub-domains of object vision (Barraclough and Perrett 2011; Kanwisher and Yovel 2006; Perrodin et al. 2015; Rossion 2014; Tsao and Livingstone 2008). Face stimuli are also convenient to use because they allow quantitative manipulation of sensory information pivotal for mechanistic characterization of the decision-making process (Waskom et al. 2019); images can be decomposed into local spatial parts (e.g., eyes, nose, mouth) and can be morphed between two instances (e.g., faces of two individuals) to create a parametric stimulus set. At the same time, human face perception is highly elaborate and embodies the central challenge of object recognition that must distinguish different identities from their complex visual appearances (Tsao and Livingstone 2008).

To quantitatively characterize the decision-making process, we investigate face recognition as a process of combining sensory evidence over both *space* and *time.* Faces are thought to be processed holistically (Maurer et al. 2002; Richler et al. 2012); breaking the configuration of facial images significantly affects face perception, indicating spatial interactions across facial parts. However, computational properties of the spatial integration remain elusive (Richler et al. 2012). One may consider that holistic recognition arises from non-linear integration of facial features (Shen and Palmeri 2015), but linear integration may also suffice to account for holistic effects (Gold et al. 2012). Furthermore, humans flexibly utilize different facial parts to categorize faces according to their behavioral needs (e.g., discrimination of identity vs. expression; Schyns et al. 2002, 2007), but the underlying mechanisms of this flexibility also remain underexplored.

Besides spatial properties, face and object recognition also include rich temporal dynamics. Although object identification and categorization are usually fast, reaction times (RTs) are often hundreds of milliseconds longer (Carlson et al. 2014; Gauthier et al. 1998; Kampf et al. 2002; Ramon et al. 2011; Witthoft et al. 2018) than the time required for a feedforward sweep along the ventral visual pathway (Hung et al. 2005; Thorpe et al. 1996). Furthermore, recognition performance follows speed-accuracy tradeoff, where additional time improves accuracy (Gauthier et al. 1997; Thorpe et al. 1996). Together, these observations suggest that the decision-making process in face and object recognition is not instantaneous but unfolds over time (Hanks et al. 2014; Heitz and Schall 2012). However, its computational properties have scarcely been characterized.

Using our novel face categorization tasks that tightly control spatiotemporal sensory information (Okazawa et al. 2021, 2018), we show that human subjects categorize faces by linearly integrating visual information over space and time. Spatial features are weighted non-uniformly and integrated largely linearly to form momentary evidence, which is then accumulated over time to generate a decision variable that guides the behavior. The temporal accumulation is also linear, and its time constant is quite long, preventing significant loss of information (leak) during the decisionmaking process. Between identity and expression categorizations, the weighting for spatial integration flexibly changes to accommodate task demands. Together, we offer a novel framework to study face recognition as a spatiotemporal integration process, which unifies two rich veins of visual research: object recognition and perceptual decision making.

## Methods

### Observers and experimental setup

Thirteen human observers (age: 18-35, 5 males and 8 females; recruited from students and staff at New York University) participated in the experiments. Observers had normal or corrected-to-normal vision. They were naïve to the purpose of the experiment, except for one observer who was an author (G.O.). They all provided informed written consent before participation. All experimental procedures were approved by the Institutional Review Board at New York University.

Throughout the experiment, subjects were seated in an adjustable chair in a semi-dark room with chin and forehead supported before a CRT display monitor (21-inch Sony GDM-5402, Tokyo, Japan; 75 Hz refresh rate; 1,600 × 1,200 pixels screen resolution; 52 cm viewing distance). Stimulus presentation was controlled with the Psychophysics Toolbox (Brainard 1997) and Matlab (Mathworks, Natick, MA). Eye movements were monitored using a high-speed infrared camera (Eyelink, SR-Research, Ontario). Gaze position was recorded at 1kHz.

### Tasks

#### Stochastic multi-feature face categorization task

The task required classification of faces into two categories, each defined by a prototype face (Fig. 1A). The subject initiated each trial by fixating a small red point at the center of the screen (FP, 0.3° diameter). After a short delay (200–500 ms, truncated exponential distribution), two targets appeared 5° above and below the FP to indicate the two possible face category choices (category 1 or 2). Simultaneously with the target onset, a face stimulus (2.18°× 2.83°) appeared on the screen parafoveally (stimulus center 1.8° to the left or right of the FP; counterbalanced across subjects; see Figs. 5-1, 5-2 for the side used for each subject). We placed the stimuli parafoveally, aiming to present the informative facial features at comparable visual eccentricities and yet keep the stimuli close enough to the fovea to take advantage of the foveal bias for face perception (Kreichman et al. 2020; Levy et al. 2001). The parafoveal presentation also enabled us to control subjects’ fixation such that small eye movements (e.g., microsaccades) within the acceptable fixation window did not substantially change the sensory inputs. Subjects reported the face category by making a saccade to one of the two targets as soon as they were ready. The stimulus was extinguished immediately after the saccade initiation. Reaction times were calculated as the time from the stimulus onset to the saccade initiation. If subjects failed to make a choice in 5 s, the trial was aborted (0.101% of trials). To manipulate task difficulty, we created a morph continuum between the two prototypes and presented intermediate morphed faces on different trials (see below). Distinct auditory feedbacks were delivered for correct and error choices. When the face was ambiguous (halfway between the two prototypes on the morph continuum), correct feedback was delivered on a random half of trials.

**Figure 1:**
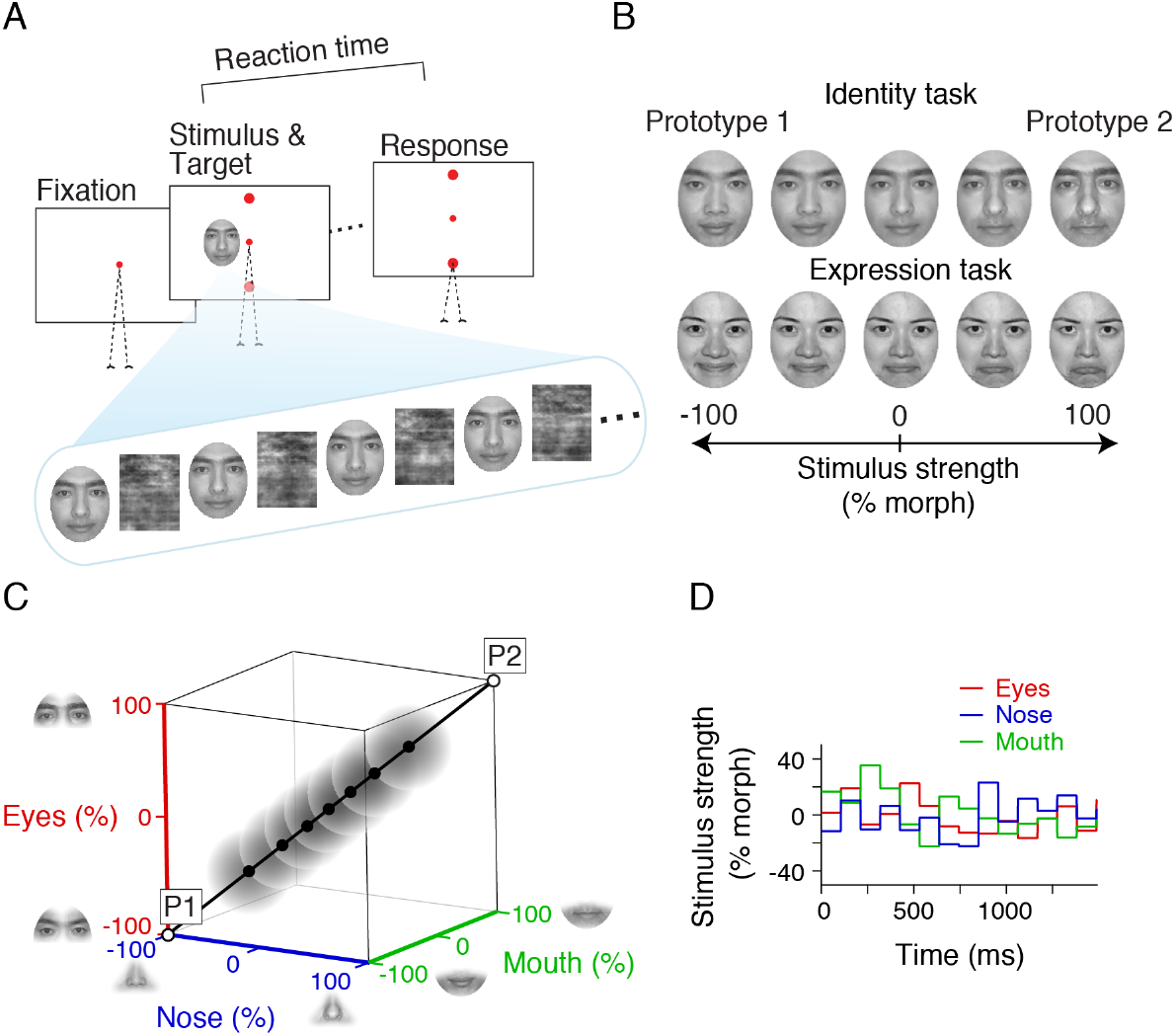
Stochastic multi-feature face categorization task. (**A**) On each trial, after the subject fixated a central fixation point, two circular targets appeared on the screen and immediately followed by a dynamic stimulus stream consisting of faces interleaved with masks. The subject was instructed to report the stimulus category (face identity or expression in different blocks) as soon as ready by making a saccadic eye movement to one of the two targets associated with the two categories. Reaction time was defined as the interval between the stimulus onset and the saccade onset. (**B**) The stimuli in each task were engineered by morphing two category prototype stimuli, defined as ±100% stimulus strengths. The prototype faces were chosen from the photographs of MacBrain Face Stimulus Set (Tottenham et al. 2009). For the identity stimuli shown in the figure, we used faces of two authors (G.O. and R.K.) to avoid copyright issues. (**C**) Our custom algorithm allowed independent morphing of different stimulus regions. We defined three regions (eyes, nose, and mouth) as informative features, while fixing other regions to the intermediate level between the prototypes. Therefore, the stimulus space for each task was three dimensional. In this space, the prototypes (P1 and P2) are two opposite corners. In each trial, the nominal mean stimulus strength was sampled from points on the diagonal line (filled black dots). Each stimulus frame was then sampled from a 3D symmetric Gaussian distribution around the specified nominal mean (gray clouds; s.d. 20% morph). (**D**) During the stimulus presentation in each trial, the morph levels of the three informative features were randomly and independently sampled from the Gaussian distribution every 106.7 ms. The features therefore furnished differential and time-varying degrees of support for the competing choices in each trial. Each face in the stimulus stream was masked with a random noise pattern created by phase scrambling images of faces (see A). The masking prevented subjects from detecting the feature changes, although they still influenced the subjects’ final choices. The stimuli were presented parafoveally to keep the informative features at more or less similar eccentricities in the visual field.

Subjects could perform two categorization tasks: identity categorization (Fig. 1B top) and expression categorization (Fig. 1B bottom). The prototype faces for each task were chosen from the photographs of MacBrain Face Stimulus Set (Tottenham et al. 2009). For the illustrations of identity stimuli in Figs. 1A-C and 1-1, we used morphed images of two authors’ faces to avoid copyright issues. We developed a custom algorithm that morphed different facial features (regions of the stimulus) independently between the two prototype faces. Our algorithm started with 97-103 manually-matched anchor points on the prototypes and morphed one face into another by linear interpolation of the positions of anchor points and textures inside the tessellated triangles defined by the anchor points. The result was a perceptually seamless transformation of the geometry and internal features from one face to another. Our method enabled us to morph different regions of the faces independently. We focused on three key regions (eyes, nose, and mouth) and created an independent series of morphs for each one of them. The faces that were used in the task were composed of different morph levels of these three informative features. Anything outside those features was set to the halfway morph between the prototypes, thus was uninformative. The informativeness of the three features (stimulus strength) was defined based on the mixture of prototypes, spanning from −100% when the feature was identical to prototype 1 to +100% when it was identical to prototype 2 (Fig. 1C). At the middle of the morph line (0% morph), the feature was equally shaped by the two prototypes.

By varying the three features independently, we could study spatial integration through creating ambiguous stimuli in which different features could support different choices (Figs. 1C, 1-1). We could also study temporal integration of features by varying the three discriminating features every 106.7ms within each trial (Fig. 1D). This frame duration allowed us sufficiently precise measurements of subjects’ temporal integration in their ~1s decision times while ensuring the smooth subliminal transition of frames (see below). The stimulus strengths of three features in each trial were drawn randomly from independent Gaussian distributions. The mean and standard deviation of these distributions were equal and fixed within each trial, but the means varied randomly from trial to trial. We tested seven mean stimulus strengths (−50%, −30%, −14%, 0%, +14%, +30%, and +50%), except for subject 13, who had a higher behavioral threshold and was tested including a higher stimulus strength (−80% and 80%). The standard deviation was 20% morph level. Sampled values that fell outside the range [−100% +100%] (0.18% of samples) were replaced with new samples inside the range. Using larger standard deviations would have allowed us to sample a wider stimulus space, but we limited the standard deviation to 20% morph level to keep the stimulus fluctuations subliminal, avoiding potential changes of decision strategy for vividly varying stimuli.

Changes in the stimulus within a trial were implemented in a subliminal fashion such that subjects did not consciously perceive variation of facial features and yet their choices were influenced by these variations. We achieved this goal using a sequence of stimuli and masks within each trial (Video 1). The stimuli were morphed faces with a particular combination of the three discriminating features. The masks were created by phase randomization (Heekeren et al. 2004) of the 0% morph face and therefore had largely matching spatial frequency content with the stimuli shown in the trial. The masks ensured that subjects did not consciously perceive minor changes in informative features over time within a trial. In debriefings following each experiment, subjects noted that they saw one face in each trial, but the face was covered with time-varying cloudy patterns (i.e., masks) over time.

For the majority of subjects (9/13), each stimulus was shown without a mask for one monitor frame (13.3ms). Then, it gradually faded out over the next seven frames as a mask stimulus faded in. For these frames, the mask and the stimulus were linearly combined, pixel-by-pixel, according to a half-cosine weighting function, such that in the last frame, the weight of the mask was 1 and the weight of the stimulus was 0. Immediately afterward, a new stimulus frame with a new combination of informative features was shown, followed by another cycle of masking, and so on. For a minority of subjects (4/13), we replaced the half cosine function for the transition of stimulus and mask with a full cosine function, where each 8-frame cycle started with a mask, transitioned to an unmasked stimulus in frame 5, and transitioned back to a full mask by the beginning of the next cycle. We did not observe any noticeable difference in the results of the two presentation methods and combined data across subjects.

Twelve subjects participated in the identity categorization task (total trials, 35,300; mean±s.d. trials per subject, 2,942±252). Seven subjects participated in the expression categorization task in separate sessions (total trials, 20,225; trials per subject, 2,889±285). Six of the subjects performed both tasks. Our subject counts are comparable to previous studies of perceptual decision-making tasks (Levi et al. 2018; Stine et al. 2020). Collecting a large number of trials from individual subjects enabled detailed quantification of decision behavior for each subject (Smith and Little 2018). Our results were highly consistent across subjects (Figs. 5-1 and 5-2). A part of the data for the identity categorization task was previously published (Okazawa et al. 2018).

#### Odd-one-out discrimination task

Our behavioral analyses and decision-making models establish that subjects’ choices in the identity and expression categorization tasks were differentially informed by the three facial features; choices were most sensitive to changes in the morph level of eyes for identity discrimination and changes in the morph level of mouth for expression discrimination (Figs. 2E-F, 5D-E). This task-dependent sensitivity to features could arise from two sources: different visual discriminability for the same features in the two tasks and/or unequal decision weights for informative features in the two tasks (Fig. 8B). To determine the relative contributions of these factors, we designed an odd-one-out discrimination task to measure visual discriminability of different morph levels of informative features in the two tasks (Fig. 8C).

**Figure 2:**
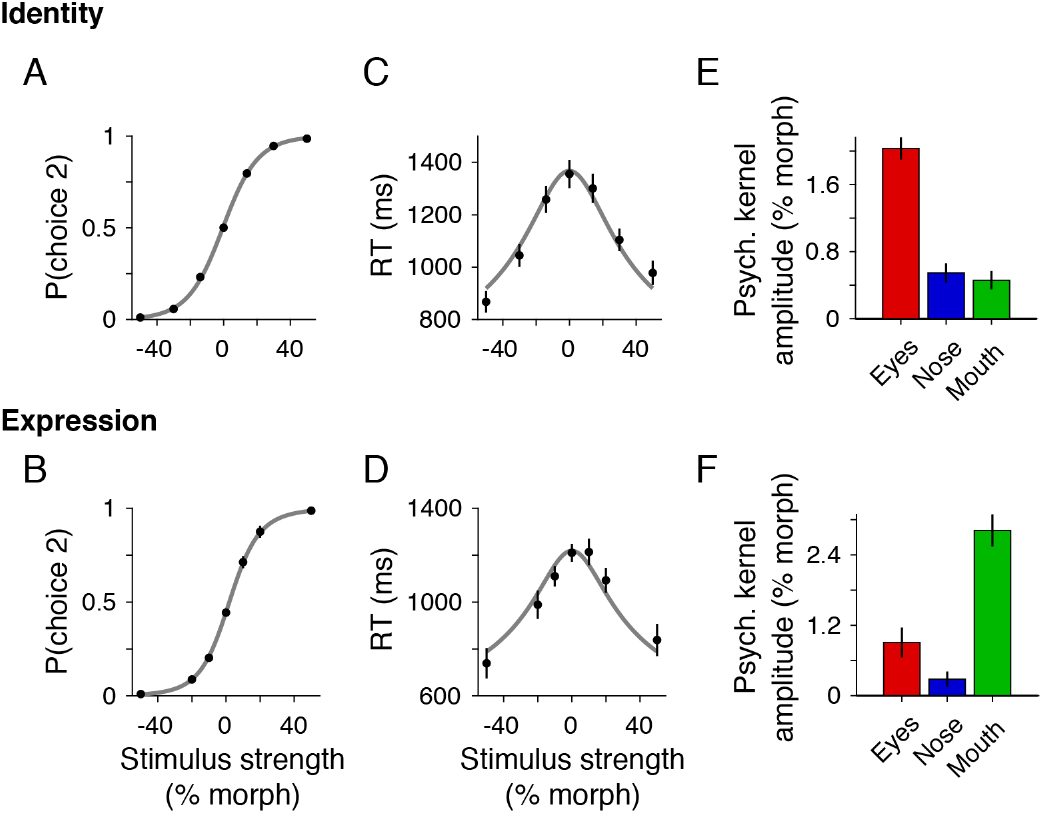
Stimulus strength shaped choices and reaction times with differential contributions from informative features. (**A, B**) Psychometric functions based on nominal stimulus strength in the identity (A) and expression (B) categorization tasks. Gray lines are logistic fits (Eq. 1). Error bars are s.e.m. across subjects (most of them buried in data dots). (**C, D**) Chronometric functions based on nominal stimulus strength in the two tasks. Average reaction times were slower for the intermediate, most ambiguous stimulus strengths. Gray lines are the fits of a hyperbolic tangent function (Eq. 2). (**E, F**) Psychophysical reverse correlation using feature fluctuations revealed positive but non-uniform contributions of multiple features in both tasks. The amplitude of the psychophysical kernel for a feature was calculated as the difference of the average feature fluctuations conditioned on the two choices. Trials with 0-14% nominal stimulus strength were used in this analysis. Error bars indicate s.e.m. across subjects. The kernel amplitudes of individual subjects are shown in Fig. 2-1. For the full time course of the kernels, see Figs. 5, 5-1, and 5-2.

On each trial, subjects viewed three stimuli presented sequentially at 1.8° eccentricity (similar to the categorization tasks). The stimuli appeared after the subject fixated a central FP and were shown for 320 ms each, with 500 ms inter-stimulus intervals. The three stimuli in a trial were the same facial feature (eyes, nose, or mouth) but had distinct morph levels, chosen randomly from the following set: {−100%, −66%, −34%, 0%, +34%, +66%, +100%}. Facial regions outside the target feature were masked by the background. The target feature varied randomly across trials. Subjects were instructed to report the odd stimulus in the sequence (the stimulus most distinct from the other two) by pressing one of the three response buttons within 2 s from the offset of the last stimulus (RT from stimulus offset, 0.66 s ± 0.13 *s,* mean±s.d.). No feedback was given after the response. Subjects underwent extensive training prior to the data collection to achieve stable and high performance. During training, two of the three stimuli were identical, and subjects received feedback on whether they correctly chose the distinct stimulus. The training continued until subjects reached 70% correct (chance level: 33%).

Nine out of the twelve subjects who participated in the identity categorization tasks also performed the odd-one-out discrimination task using identity stimuli in separate blocks of the same sessions. Three out of the seven subjects who participated in the expression tasks performed the odd-one-out task using expression stimuli. For the identity stimuli,13,648 trials were collected across the three features (9 subjects; 1,516±420 trials per subject, mean±s.d.). For the expression stimuli, 3,570 trials were collected (3 subjects; 1,190±121 trials per subject).

### Data analysis

#### Psychometric and chronometric functions

We assessed the effects of stimulus strength on the subject’s performance by using logistic regression (Fig. 2A-B):

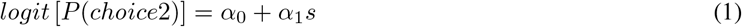

where *logit*(*p*) = *log*(*p*/1 – *p*), s is the nominal stimulus strength ranging from −1 (−100% morph level) to +1 (+100% morph level), and *α_i_* are regression coefficients. *α*_0_ quantifies the choice bias and α_1_ quantifies the slope of the psychometric function.

The relationship between the stimulus strength and the subject’s mean reaction times (RTs) was assessed using a hyperbolic tangent function (Shadlen et al. 2006; Fig. 2C-D):

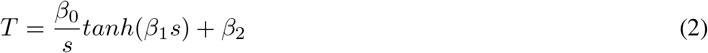

where T is the mean RTs measured in milliseconds and *β_i_* are model parameters. *β*_0_ and *β*_1_ determine the stimulusdependent changes in decision time, whereas *β*_2_ quantifies the sensory and motor delays that elongate the RTs but are independent of the decision-making process (i.e., non-decision time).

#### Psychophysical reverse correlation

To quantify the effect of stimulus fluctuations over time and space (facial features) on choice (Fig. 1D), we performed psychophysical reverse correlation (Ahumada 1996; Okazawa et al. 2018) (Figs. 2E-F, 5D-E, 7D-E). Psychophysical kernels (*K_f_*(*t*)) were calculated as the difference of average fluctuations of morph levels conditional on the subject’s choices:

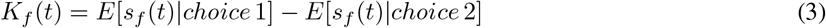

where *s_f_*(*t*) is the morph level of feature *f* at time *t*. This analysis only used trials with low stimulus strength (nominal morph level, 0-14%; 14,213 trials across twelve subjects in the identity task and 7,882 trials across seven subjects in the expression task). For the non-zero strength trials, the mean strength was subtracted from the fluctuations, and the residuals were used for the reverse correlation. Figs. 5D-E and 7D-E show the time course of psychophysical kernels for the three informative features. We use stimulus fluctuations up to the median RT aligned to stimulus onset or the saccade onset in order to ensure that at least half of the trials contributed to the kernels at each time. Fig. 2E-F shows the kernels averaged over time from stimulus onset to median RT. For the psychophysical kernels shown in Figs. 2E-F, 5D-E, 7D-E, we aggregated data across all subjects. The kernels of individual subjects are shown in Figs. 5-1 and 5-2. We did not perform any smoothing on the kernels.

#### Joint psychometric function

To quantify the effect that co-fluctuations of feature strengths have on choice, we quantified the probability of choices as a function of the joint distribution of the stimulus strengths across trials (Fig. 3A-B). We constructed the joint distribution of the three features by calculating the average strength of each feature in the trial. Thus, one trial corresponds to a point in a 3D feature space (see Fig. 1C). In this space, the probability of choice was computed within a Gaussian window with a standard deviation of 4%. Fig. 3A-B shows 2D intersections of this 3D space. We visualized the probability of choice by drawing iso-probability contours at 0.1 intervals. The trials of all stimulus strengths were included in this analysis, but similar results were also obtained by restricting the analysis to the low morph levels (≤14%). We aggregated data across all subjects, but similar results were observed within individual subjects.

**Figure 3:**
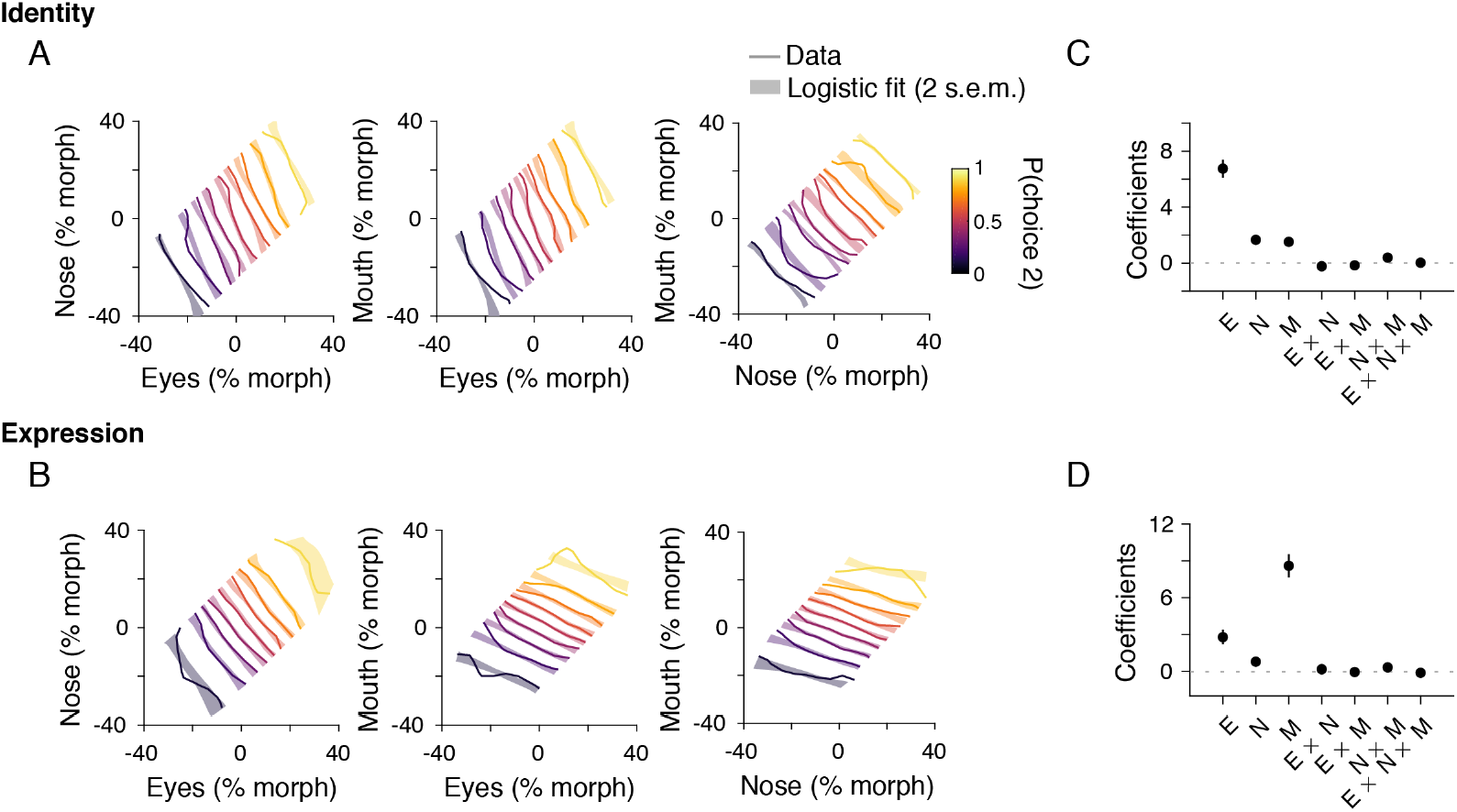
Spatial integration across informative features was largely linear. (**A, B**) Two-dimensional slices of the psychometric function for the three-dimensional stimulus space. Each panel shows the iso-probability contours for choice 2 as a function of the true average morph level of two of the three informative features in each trial. The third informative feature was marginalized for making each panel. The iso-probability contours are drawn at 0.1 intervals and moderately smoothed with a 2D Gaussian function (s.d., 4% morph). Thin lines are actual data, and thick pale lines are logistic fit (Eq. 4; the thickness of the lines reflects 2 s.e.m. of the fitted parameters). The straight and parallel contour lines are compatible with a largely linear spatial integration process across features, with the slope of the contours reflecting the relative contribution of the features illustrated in each panel. (**C, D**) A logistic regression to evaluate the relative contribution of individual features and their multiplicative interactions to subjects’ choices supported a largely linear spatial integration process: the interaction terms have minimal impact on choice beyond the linear integration across features. E, N, and M indicate eyes, nose, and mouth, respectively. Error bars indicate s.e.m. across subjects.

To quantify linear and multiplicative effects on joint psychometric functions, we performed the following logistic regression:

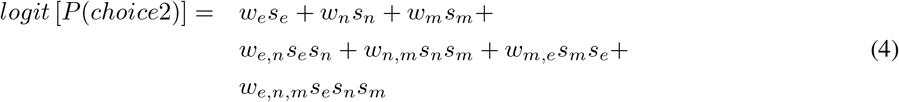

where *s_e_, s_n_*, and *s_m_* corresponds to the stimulus strengths of eyes, nose, and mouth for individual stimulus frames. *w_e_*, *w_n_*, and *w_m_* are model coefficients for linear factors, whereas *w_e,n_, w_n,m_, w_m,e_,* and *w_e,n,m_* are coefficients for multiplicative factors. In this regression, the dynamic ranges of the linear and multiplicative terms were scaled to match in order to ensure more homogeneous distribution of explainable variance across different factors; otherwise, the fluctuations of multiplicative terms would be one or two orders of magnitude smaller than those of the linear terms. Figure 3C-D shows the coefficients averaged across subjects. The results from individual subjects are shown in Fig. 3-1.

#### Relationship between stimulus strength and subjective evidence

To quantitatively predict behavioral responses from stimulus parameters, one must first know the mapping function between the physical stimulus strength (morph level) and the amount of evidence subjects acquired from the stimulus. This mapping could be assessed by performing a logistic regression that relates choice to different ranges of stimulus strength (Fig. 4), similar to those performed in previous studies (Waskom and Kiani 2018; Yang and Shadlen 2007). For this analysis, we used the following regression:

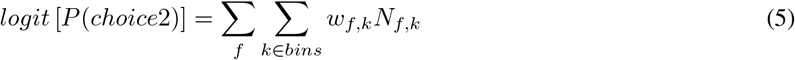

where *N_f,k_* is the number of stimulus frames that fall into decile *k* for feature *f* in a given trial, and *w_f,k_* are regression coefficients that signify the subjective evidence assigned to a morph level *k* of feature *f*. Division of feature morph levels into deciles in our regression aimed at limiting the number of free parameters while maintaining adequate resolution to quantify the mapping function between the stimulus strength and momentary evidence. If the subjective evidence in units of log-odds scales linearly with the morph level, then *w_f,k_* would linearly change with the morph level. Plotting *w_f,k_* as a function of feature morph level indicated a linear relationship, except perhaps for the extreme deciles at the two ends of the morph line (Fig. 4). For illustration purposes, the fitting lines in Figure 4 exclude the extreme deciles, but to ensure unbiased reporting of statistics, we included all the deciles to quantify the accuracy of the linear fit in Results.

**Figure 4:**
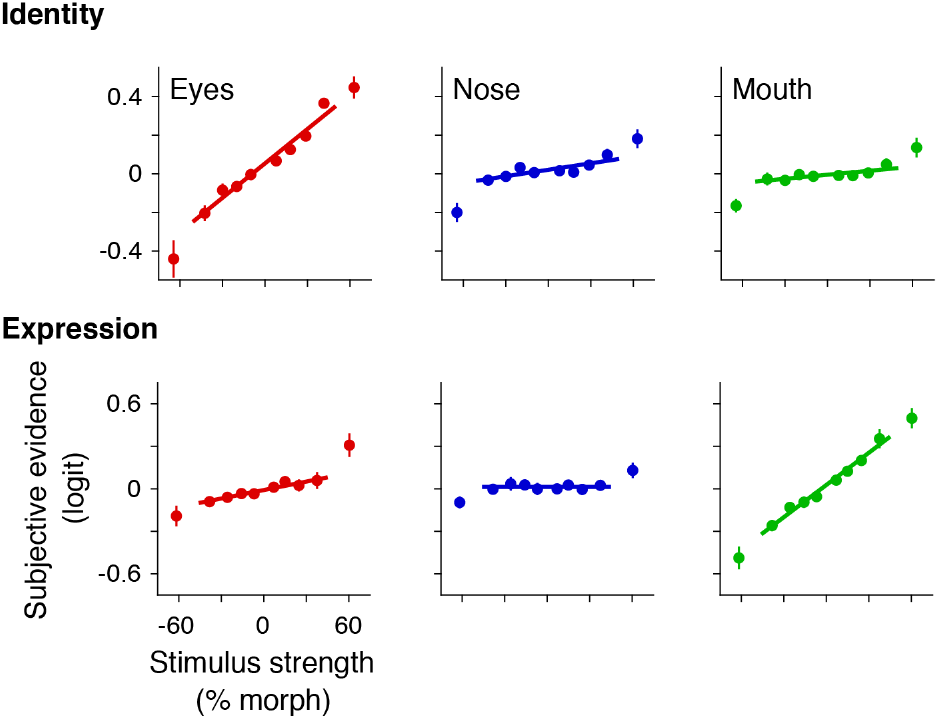
The evidence conferred by a feature mapped linearly to the feature morph level, especially for the intermediate stimuli. The linear integration of features across space allowed accurate quantification of the evidence that subjects inferred from each feature. We split morph levels of each feature into ten quantiles and used a logistic regression that explained choices based on the number of occurrences of the quantiles in each trial, quantifying the subjective evidence of each morph level in units of log odds of choice. Subjective evidence linearly scaled with the morph level for each feature, except for the highest strengths. The lines are linear regressions of subjective evidence against feature morphs, excluding the highest strengths. Error bars indicate s.e.m. across subjects.

### Model fit and evaluation

To quantitatively examine the properties of the decision-making process, we fit several competing models to the subject’s choices and RTs. Based on our earlier analyses (Figs. 3 and 4), these models commonly use a linear mapping between feature morph levels and the evidence acquired from each feature, as well as linear functions for spatial integration of informative features in each frame. The combined momentary evidence from each stimulus frame was then integrated over time. Our main models are therefore extensions of the drift diffusion model, where fluctuations of the three informative facial features are accumulated toward decision bounds and reaching a bound triggers a response after a non-decision time. Our simplest model used linear integration over time, whereas our more complex alternatives allowed leaky integration or dynamic changes of sensitivity over time. We also examined models without spatial integration, where the evidence from each informative feature was accumulated independently (i.e., three competing drift diffusion processes) and the decision and RT were determined by the first process reaching a bound. Below, we first provide the equations and intuitions for the simplest model and explain our fitting and evaluation procedures. Afterward, we explain the alternative models.

Spatial integration in our models linearly combines the strength of features at each time to calculate the momentary evidence conferred by a stimulus frame:

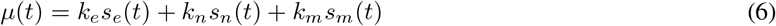

where *s_e_*(*t*), *s_n_*(*t*), *s_m_*(*t*) the strengths of eyes, nose, and mouth at time *t*, and *k_e_, k_n_, k_m_* are the sensitivity parameters for each feature. Momentary evidence (*μ*(*t*)) was integrated over time to derive the decision variable. The process stopped when the decision variable reached a positive or negative bound (±B). The probability of crossing the upper and lower bounds at each decision time can be calculated by solving the Fokker-Planck equation (Karlin and Taylor 1981; Kiani and Shadlen 2009):

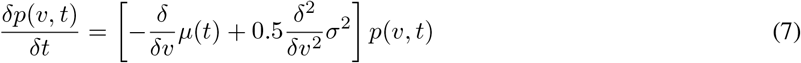

where *p*(*v, t*) is the probability density of the decision variable at different times. The boundary conditions are

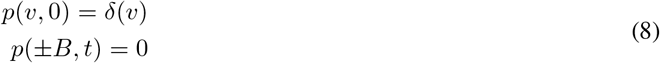

where *δ*(*v*) denotes a delta function. The first condition enforces that the decision variable always starts at 0, and the second condition guarantees that the accumulation terminates when the decision variable reaches one of the bounds. We set the diffusion noise (*σ*) to 1 and defined the bound height and drift rate in units of σ. RT distribution for each choice was obtained by convolving the distribution of bound crossing times with the distribution of non-decision time, which was defined as a Gaussian distribution with a mean of *T*_0_ and standard deviation of *σ*_*T*0_.

Overall, this linear integration model had six degrees of freedom: decision bound height (*B*), sensitivity parameters (*k_e_, k_n_, k_m_*), and the parameters for non-decision time (*T*_0_, *σ*_*T*0_). We fit model parameters by maximizing the likelihood of the joint distribution of the observed choices and RTs in the experiment (Okazawa et al. 2018). For a set of parameters, the model predicted the distribution of RTs for each possible choice for the stimulus strengths used in each trial. These distributions were used to calculate the log-likelihood of the observed choice and RT on individual trials. These log-likelihoods were summed across trials to calculate the likelihood function for the dataset. Model parameters were optimized to maximize this function. To avoid local maxima, we repeated the fits from 10 random initial points and chose the fit with the highest likelihood. The trials of all stimulus strengths were included in this fitting procedure. The fits were performed separately for each subject. Figure 5B, C shows the average fits across subjects.

**Figure 5:**
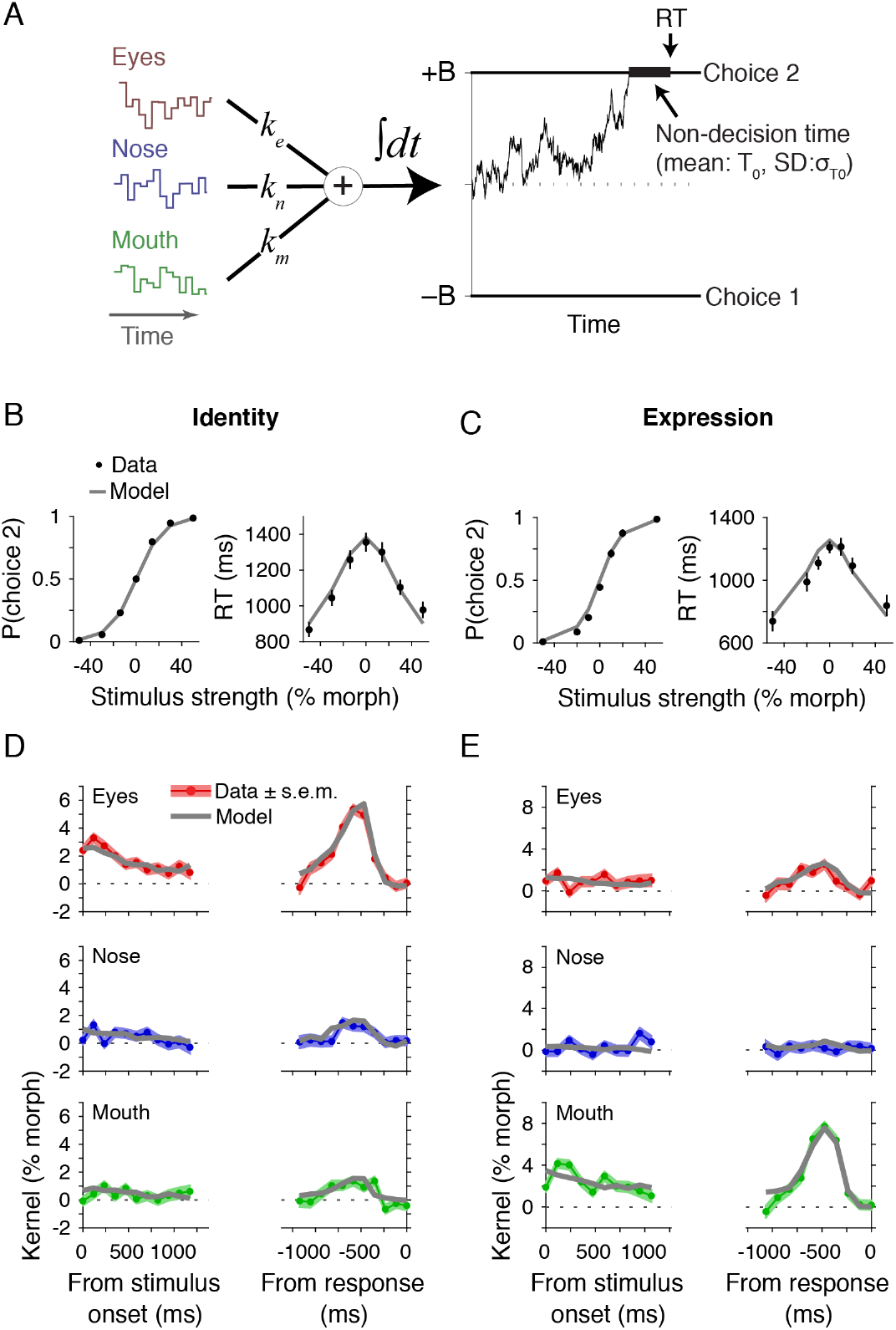
A model that linearly integrates sensory evidence over space and time accounted for choice, reaction time, and psychophysical kernels. (**A**) Multi-feature drift diffusion model. The model linearly integrates the morph levels of the three informative features with static spatial sensitivities (*k_e_, k_n_, k_m_*) to create the momentary evidence, which is then integrated over time to create a decision variable. The integration process continues until the decision variable reaches one of the two decision bounds, corresponding to choices 1 and 2. Reaction time equals the time to reach the bound (decision time) plus a non-decision time that reflects the aggregate of sensory and motor delays. (**B, C**) Model fit to choices and reaction times in the identity (B) and expression (C) categorization tasks. Data points (dots) are the same as those in Fig. 2A-D. (**D, E**) The model also accurately explained the amplitude and dynamics of psychophysical kernels. Shading indicates s.e.m. The differences across tasks are due to distinct feature sensitivities.

To generate the model psychophysical kernels, we created 10^5^ test trials with 0% stimulus strength using the same stimulus distributions as in the main task (i.e., Gaussian distribution with a standard deviation of 20%). We simulated model responses for these trials with the same parameters fitted for each subject. We then used the simulated choices and RTs to calculate the model prediction for psychophysical kernels of the three features. Figure 5D-E shows the average predictions superimposed on the observed psychophysical kernels, averaged across subjects (for single subject comparisons see Figs. 5-1 and 5-2). Note that the model kernels were not directly fit to match the data. They were calculated based on an independent set of simulated 0% stimulus trials, making the comparison in Fig. 5D, E informative.

The same fitting procedure was used for the alternative models explained below.

#### Leaky integration

To test the degree of temporal integration, we added a memory loss (leak) in the decision-making process. This model is implemented as an Ornstein-Uhlenbeck process, whose Fokker-Planck equation is:

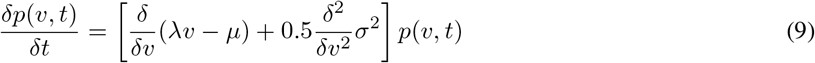

where λ is the leak rate. A larger leak rate indicates greater loss of information over time. At the limit of infinitely large leak rate, the model no longer integrates evidence and makes decisions based solely on whether the most recently acquired momentary evidence exceeds one of the decision bounds.

#### Dynamic sensitivity

To test whether the effect of sensory evidence on choice is constant over time, we allowed sensitivity to features to be modulated dynamically. To capture both linear and non-linear temporal changes, the modulation included linear (*γ*_1_) and quadratic (*γ*_2_) terms such that

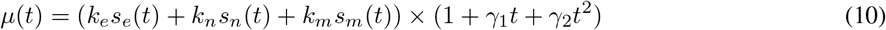

#### Parallel accumulation of evidence from three facial features

The models above first integrated the evidence conferred by the three informative facial features (spatial integration) and then accumulated this aggregated momentary evidence over time. We also considered alternative models, in which evidence from each feature was accumulated independently over time. These models therefore included three competing accumulators. Each accumulator received momentary evidence from one feature with fixed sensitivity (*k_e_, k_n_, k_m_* for eyes, nose, mouth). In the model featured in Fig. 7A, the accumulator that first reached a decision bound dictated the choice and decision time. As in the models above, a non-decision time separated the bound crossing and response time.

To further explore different decision rules, we constructed two variants of the parallel accumulation model (Fig. 7-1). In the first variant, the decision was based on the sign of the majority of the accumulators (i.e., two or more out of three accumulators) at the moment when one accumulator reached its bound. In the second variant, the decision was based on the sign of the sum of the decision variables across the three accumulators at the time when one accumulator reached its bound. All model variants had six free parameters (*k_e_, k_n_, k_m_, B, T*_0_, *σ*_*T*0_), equal to the degrees of freedom of the main model explained above.

### Analysis of odd-one-out discrimination task

We used subjects’ choices in the odd-one-out task to estimate visual discriminability of different morph levels of the informative features. We adopted an ideal observer model developed by Maloney and Yang (2003), where the perception of morph level *i* of feature *f* is defined as a Gaussian distribution, 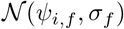, with mean *ψ_i,f_* and s.d. *σ_f_*. The discriminability of a triad of stimuli is determined by their perceptual distances (|*ψ_i,j_* – *ψ*_2,*f*_|, |*ψ*_2,*f*_ – *ψ*_3,*f*_|, |*ψ*_3,*f*_ – *ψ*_1,*f*_|). Specifically, an ideal observer performing the odd-one-out discrimination of three morph levels (*i, j, k*) of feature *f* would choose *i* if the perceptual distances of *i* from *j* and k are larger than the perceptual distance of *j* and *k*:

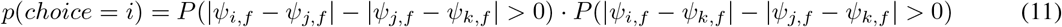

The probabilities on the right side of the equation can be derived from 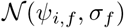. The probability of choosing *j* and *k* can be calculated in a similar way (Maloney and Yang 2003). The model captures both the discriminability of morph levels of the same feature, through *ψ_i,f_*, and differences across features, through *σ_f_*.

We fit the ideal observer model to the subject’s choices using maximum likelihood estimation. Since there were seven morph levels for each feature in our task, choices for each feature could be explained using eight parameters (*ψ_i,f_* for *i* = 1... 7, and *σ_f_*). Six of these parameters are free (*ψ*_2,*f*_ ⋯ *ψ*_6,*f*_ and *σ_f_*). *ψ*_1,*f*_ and *ψ*_7,*f*_ were anchored at −1 and +1, respectively, to avoid redundant degrees of freedom in the fits. The model was fit separately for each subject and feature. To avoid local maxima, we repeated the fits from 10 random initial points and chose the parameters that maximized the likelihood function. Because *ψ_i,f_* changed largely linearly with the feature strength (% morph) in all fits (Fig. 8-2A) and the range of *ψ_i,f_* was fixed at [-1, +1], we could quantify perceptual discriminability of different features using their respective *σ_f_*. Specifically, *d*′ between the two extreme morph levels of a feature is 2/*σ_j_* as

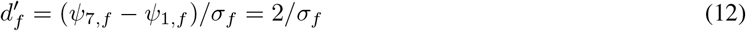

If the differences of feature sensitivity parameters (*k_e_, k_n_, k_m_* in Eq. 6) in the categorization tasks were fully determined by the visual discriminability of features – i.e., there was no task-dependent weighting of the features – *k_f_* would have the same relative scales as the 1/*σ_f_*. To test this, we divided the model sensitivities for the three facial features by their respective 1/*σ_f_* to estimate task-dependent decision weights. The resulting decision weights showed significant inhomogeneity across features and between tasks (Fig. 8F), suggesting the presence of task-dependent weighting of facial features categorization.

## Results

### Spatial integration in face categorization

We developed stochastic multi-feature face categorization tasks suitable for studying spatial and temporal properties of the computations underlying the decision-making process. Subjects classified naturalistic face stimuli into two categories. In each trial, subjects observed a face stimulus with subliminally varying features and, when ready, reported its category with a saccadic eye movement to one of the two targets (Fig. 1A). The targets were associated with the two prototypes that represented the discriminated categories: identities of two different persons in the identity categorization task (Fig. 1B top) or happy and sad expressions of a person in the expression categorization task (Fig. 1B bottom). The stimulus changed dynamically in each trial. The dynamic stimulus stream consisted of a sequence of face stimuli interleaved by masks (Fig. 1A). Each face stimulus was engineered to have three informative features in the eyes, nose, and mouth regions, and sensory evidence conferred by the three informative features rapidly fluctuated over time as explained in the next paragraph. The masks between face stimuli kept the changes in facial features subliminal, creating the impression that a fixed face was covered periodically with varying noise patterns (see Methods; Video 1).

Using a custom algorithm, we could independently morph the informative facial features (eyes, nose, mouth) between the two prototypes to create a three-dimensional (3D) stimulus space whose axes correspond to the morph level of the informative features (Figs. 1C, 1-1). In this space, the prototypes are two opposite corners (specified as ±100% morph), and the diagonal connecting the prototypes correspond to a continuum of faces where the three features of each face equally support one category vs. the other. For the off-diagonal faces, however, the three features provide unequal or even opposing information for the subject’s choice. In each trial, the nominal mean stimulus strength (% morph) was sampled from the diagonal line (black dots in Fig. 1C). The dynamic stimulus stream was created by independently sampling a stimulus every 106.7ms from a 3D symmetric Gaussian distribution with the specified nominal mean (s.d. 20% morph; Fig. 1C-D). The presented stimuli were therefore frequently off-diagonal in the stimulus space. The subtle fluctuations of features influenced subjects’ choices as we show below, enabling us to determine how subjects combined spatiotemporal sensory evidence over space and time for face categorization.

We first evaluated subjects’ choices and RTs in both tasks (Fig. 2). The average correct rate excluding 0% morph level was 91.0% ± 0.7% (mean±s.e.m. across subjects) for the identity task and 89.2% ±1.2% for the expression task. The choice accuracy monotonically improved as a function of the nominal mean stimulus strength in the trial (Fig.2A, B; identity task, *α*_1_ = 9.6 ± 1.8 in Eq. 1, *t*(11) = 18.3, *p* = 1.4 × 10^−9^; expression task, *α*_1_ = 11.3 ± 3.0, *t*(6) = 9.8, *p* = 6.4 × 10^−5^, two-tailed t-test). Correspondingly, the reaction times became faster for higher stimulus strengths (Fig. 2C, D; identity task, *β*_1_ = 4.7 ± 1.0 in Eq. 2, *t*(11) = 16.4, *p* = 4.4 × 10^−9^; expression task, *β*_1_ = 4.9 ± 1.9, *t*(6) = 6.6, *p* = 5.6 × 10^−4^). These patterns are consistent with evidence accumulation mechanisms that govern perceptual decisions with simpler stimuli, e.g., direction discrimination of random dots motions (Palmer et al. 2005; Ratcliff and Rouder 2000; Smith and Vickers 1988). However, the decision-making mechanisms that do not integrate sensory evidence over time can also generate qualitatively similar response patterns (Stine et al. 2020; Waskom and Kiani 2018). Furthermore, because in our task design, we used identical nominal mean morph levels for the informative features in a trial, characterizing behavior based on the mean levels can not reveal if subjects integrated sensory evidence across facial features (spatial integration). However, we can leverage the stochastic fluctuations of the stimulus to test if sensory evidence was integrated over space and time. In what follows, we first quantify the properties of spatial integration and then examine the properties of temporal integration.

To test if multiple facial features informed subjects’ decisions, we used psychophysical reverse correlation to evaluate the effect of the fluctuations of individual features on choice. Psychophysical kernels were generated by calculating the difference between the feature fluctuations conditioned on choice (Eq. 3). We focused on trials with the lowest stimulus strengths where choices were most strongly influenced by the feature fluctuations (0-14%; mean morph level of each trial was subtracted from the fluctuations; see Methods). Fig. 2E, F shows the kernel amplitude of the three facial features averaged over time from stimulus onset to median RT. These kernel amplitudes quantify the overall sensitivity of subjects’ choices to equal fluctuations of the three features (in %morph units). The kernel amplitudes markedly differed across features in each task (identity task: *F*(2, 33) = 55.4, *p* = 2.8 × 10^−11^, expression task: *F*(2,18) = 33.6, *p* = 8.5 × 10^−7^; one-way ANOVA), greatest for the eyes region in the identity task (*p* < 9.5 × 10^−10^ compared to nose and mouth, post-hoc Bonferroni test) and for the mouth region in the expression task (*p* < 3.9 × 10^−5^ compared to eyes and nose).

Critically, the choice was influenced by more than one feature. In the identity task, all three features had significantly positive kernel amplitudes (eyes; *t*(11) = 15.4, *p* = 8.6 × 10^−9^, nose: *t*(11) = 4.8, *p* = 5.4 × 10^−4^, mouth: *t*(11) = 4.2, *p* = 0.0015, two-tailed t-test for each feature). In the expression task, mouth and eyes had statistically significant kernel amplitudes and nose had a positive kernel, although it did not reach significance (mouth: *t*(6) = 10.3, *p* = 4.8 × 10^−5^, eyes: *t*(6) = 3.6, *p* = 0.012, nose: *t*(6) = 2.2, *p* = 0.072). Positive kernels for multiple facial features were prevalently observed for individual subjects too (Fig. 2-1). Therefore, the pooled results are not due to mixing data from multiple subjects with distinct behavior. These results suggest that subjects use multiple facial features for categorization, but the features non-uniformly contribute to their decisions, and their relative contributions differ between the tasks (interaction between feature kernels and tasks: *F*(2,10) = 90.5, *p* = 3.9 × 10^−7^; two-way repeated-measures ANOVA with 6 subjects who performed both tasks) — a topic we will revisit in the following sections.

Although the amplitude of psychophysical kernels informs us about the overall sensitivity of choice to feature fluctuations in the face stimuli, it does not clarify the contribution of sensory and decision-making processes to this sensitivity. Specifically, subjects’ choices may be more sensitive to changes in one feature because the visual system is better at discriminating the feature changes (visual discriminability) or because the decision-making process attributes a larger weight to the changes of that feature (decision weights) (Schyns et al. 2002; Sigala and Logothetis 2002). We will dissociate these factors in the last section of Results, but for the intervening sections, we will focus on the overall sensitivity of choice to different features.

### Linearity of spatial integration of facial features

How do subjects integrate information from multiple spatial features? Could it be approximated as a linear process, or does it involve significant non-linear effects —e.g., synergistic interactions that magnify the effect of co-fluctuations across features? Non-linear effects can be empirically discovered by plotting joint psychometric functions that depict subjects’ accuracy as a function of the strength of the facial features (Fig. 3A, B). Here, we define the true mean strength of each feature as the average of the feature morph levels over the stimulus frames shown on each trial (for temporal effects, see Figs. 5 and 6). The plots visualize the three orthogonal two-dimensional (2D) slices of the 3D stimulus space (Fig. 1C), and the contour lines show the probability of choosing the second target (choice 2) at the end of the trial.

**Figure 6:**
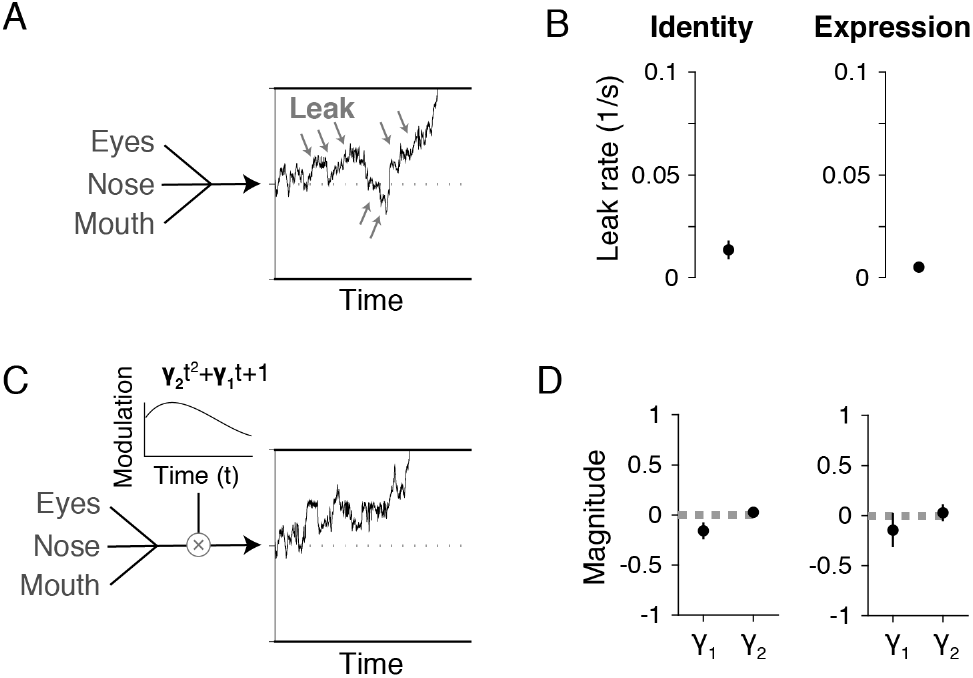
Spatiotemporal integration had static sensitivity to features and minimal forgetting (leak). We used two extensions of the multi-feature drift diffusion model to test for leaky integration and modulation of feature sensitivities over time. (**A**) Schematic of the leaky integration model. An exponential leak term was added to the temporal integration process to examine potential loss of information over time (Eq. 9). The leak pushes the decision variable toward zero (gray arrows), reducing the effect of earlier evidence. (**B**) Fitting the leaky integration model to behavioral data revealed a leak rate close to zero for both the identity and expression categorization tasks. (**C**) Schematic of the model with dynamic modulation of feature sensitivities. We added a second-order polynomial modulation function to investigate potential temporally non-uniform influence of stimulus features on choice within a trial (Eq. 10). The function allows for a variety of temporal patterns, including ascending, descending, and non-monotonic changes. (**D**) Fitting the model with dynamic sensitivities revealed no substantial change over time, with both the linear and quadratic terms of the modulation function close to zero. Error bars indicate s.e.m. across subjects.

These iso-performance contours (thin lines in Fig. 3A, B) were largely straight and parallel to each other, suggesting that a weighted linear integration across features underlies behavioral responses. The slope of contours in each 2D plot reflects the relative contribution of the two facial features to choice. For example, the nearly vertical contours in the eyes vs. nose plot of the identity task indicate that eyes had a much greater influence on subjects’ choices, consistent with the amplitudes of psychophysical kernels (Fig. 2E). Critically, the straight and parallel contour lines indicate that spatial integration does not involve substantial non-linearity. A linear model, however, does not explain curved contours, which appear at the highest morph levels, especially in the 2D plots of the less informative pairs (e.g., the Nose x Mouth plot for the identity task). Multiple factors could give rise to the curved contours. First, subjects rarely make mistakes at the highest morph levels, reducing our ability to perfectly define the contour lines at those levels. Second, the 2D plots marginalize over the third informative feature and this marginalization is imperfect due to finite trial counts in the dataset. Put together, we cannot readily attribute the presence of curved contours at the highest morph levels to non-linear processes and should rely on statistical tests for their discovery. As we explain below, statistical tests fail to detect non-linearity in the integration of features.

To quantify the contributions of linear and non-linear factors, we performed a logistic regression on the choices using both linear and non-linear multiplicative combinations of the feature strengths (Eq. 4). The model accurately fit to the contour lines in Fig. 3A, B (thick pale lines in Fig. 3A, B; identity task, *R*^2^ = 0.998; expression task, *R*^2^ = 0.999; the thickness of the lines reflects 2 s.e.m. of the logistic parameters). The model coefficients (Fig. 3C, D) show significant positive sensitivities for the linear effects of all facial features in the identity task (eyes: *t*(11) = 10.5, *p* = 4.3 × 10^−7^, nose: *t*(11) = 4.6, *p* = 8.2 × 10^−4^, mouth: *t*(11) = 4.8, *p* = 5.2 × 10^−4^, two-tailed t-test) and in the expression task (eyes: *t*(6) = 4.7, *p* = 0.0032, nose: *t*(6) = 3.4, *p* = 0.015, mouth: *t*(6) = 9.2, *p* = 9.5 × 10^−5^), but no significant effect for non-linear terms (*p* > 0.27 for all multiplicative terms in both tasks). These results were largely consistent for individual subjects, too (Fig. 3-1; 11/12 subjects in identity and 7/7 in expression task showed no significant improvement in fitting performance by adding non-linear terms (*p* > 0.05); likelihood ratio test, Bonferroni corrected across subjects). Overall, linear integration provides an accurate and parsimonious account of how features were combined over space for face categorization.

### Linearity of the mapping between stimulus strength and subjective evidence

Quantitative prediction of behavior requires understanding the mapping between the stimulus strength as defined by the experimenter (morph level in our experiment) and the evidence conferred by the stimulus for the subject’s decision. The parallel linear contours in Fig. 3 demonstrate that the strength of one informative feature can be traded for another informative feature to maintain the same choice probability. They further show that this tradeoff is largely stable across the stimulus space, strongly suggesting a linear mapping between morph levels and inferred evidence.

To formally test this hypothesis, we quantified the relationship between feature strengths and their effects on choice by estimating subjective evidence in log-odds units. Following the methods developed by Yang et al. (2007), we split the feature strengths (% morph) of each stimulus frame into ten levels and performed a logistic regression to explain subjects’ choices based on the number of occurrences of different feature morph levels in a trial. The resulting regression coefficients correspond to the change of the log-odds of choice furnished by a feature morph level. For both the identity and expression morphs, the stimulus strength mapped linearly onto subjective evidence (Fig. 4; identity task: *R*^2^ = 0.94, expression task: *R*^2^ = 0.96), with the exception of the highest stimulus strengths, which exerted slightly larger effects on choice than expected from a linear model. The linearity for a wide range of morph levels —especially for the middle range in which subjects frequently chose both targets— permits us to approximate the total evidence conferred by a stimulus as a weighted sum of the morph levels of its informative features.

### Temporal integration mechanisms

The linearity of spatial integration significantly simplifies our approach to investigate integration of sensory evidence over time. We adopted a quantitative model-based approach by testing a variety of models that have the same linear spatial integration process but differ in ways that use stimulus information over time. We leveraged stimulus fluctuations within and across trials to identify the mechanisms that shaped the behavior. We further validated these models by comparing their predicted psychophysical kernels with the empirical ones.

In our main model, the momentary evidence from each stimulus frame is linearly integrated over time (Fig. 5A). The momentary evidence from a stimulus frame is a linear weighted sum of the morph levels of informative features in the stimulus, compatible with linear spatial integration shown in the previous sections. The model assumes that sensitivities for these informative features (*k_e_, k_n_, k_m_*) are fixed within a trial. However, because the stimulus is dynamic and stochastic, the rate of increase of accumulated evidence varies over time. The decision-making process is noisy, with the majority of noise originating from the stimulus representation in sensory cortices and inference of momentary evidence from these representations (Brunton et al. 2013; Drugowitsch et al. 2016; Waskom and Kiani 2018). Due to the noise, the decision-making process resembles a diffusion process with variable drift over time. The process stops when the decision variable (accumulated noisy evidence) reaches one of the two bounds, corresponding to the two competing choices in the task. The bound that is reached dictates the choice, and the reaction time equals the time to bound (decision time) plus a non-decision time composed of sensory and motor delays (Gold and Shadlen 2007; Link 1992; Ratcliff and Rouder 2000; Smith and Vickers 1988). For each stimulus sequence, we calculated the probability of different choices and expected distribution of reaction times, adjusting model parameters to best match the model with the observed behavior (maximum likelihood fitting of the joint distribution of choice and RT; see Methods). The model accurately explained subjects’ choices (identity: *R*^2^ = 0.99 ± 0.002, expression: *R*^2^ = 0.98 ± 0.005, mean±s.e.m. across subjects) and mean reaction times (identity: *R*^2^ = 0.86 ± 0.05, expression: *R*^2^ = 0.81 ± 0.05) (Fig. 5B, C), as well as the distributions of reaction times (Fig. 5-3). The accurate match between the data and the model was also evident within individual subjects (Figs. 5-1 and 5-2).

The same model also quantitatively explains the psychophysical kernels (Fig. 5D, E; identity task: *R*^2^ = 0.86, expression task: *R*^2^ = 0.84). The observed kernels showed characteristic temporal dynamics besides the inhomogeneity of amplitudes across features, as described earlier (Fig. 2E, F). The temporal dynamics are explained by decision bounds and non-decision time in the model (Okazawa et al. 2018). When aligned to stimulus onset, the kernels decreased over time. This decline happens in the model because non-decision time creates a temporal gap between bound crossing and the report of the decision, making stimuli immediately before the report inconsequential for the decision. When aligned to the saccadic response, the kernels peaked several hundred milliseconds prior to the saccade. This peak emerges in the model because stopping is conditional on a stimulus fluctuation that takes the decision variable beyond the bound, while the drop near the response time happens again because of the non-decision time. Critically, the model assumptions about static sensitivity and linear integration matched the observed kernels. Further, the inequality of kernel amplitudes across facial features and tasks were adequately captured by the different sensitivity parameters for individual features (*k_e_, k_n_, k_m_*) in the model.

To further test properties of temporal integration, we challenged our model with two plausible extensions (Fig. 6). First, integration may be imperfect, and early information can be gradually lost over time (Bogacz et al. 2006; Usher and McClelland 2001). Such a leaky integration process can be modeled by incorporating an exponential leak rate in the integration process (Fig. 6A). When this leak rate becomes infinitely large, the model reduces to a memoryless process that commits to a choice if the momentary sensory evidence exceeds a decision bound — i.e., extrema detection (Stine et al. 2020; Waskom and Kiani 2018). To examine these alternatives, we fit the leaky integration model to the behavioral data. Although the leak rate is difficult to assess in typical perceptual tasks (Stine et al. 2020), our temporally fluctuating stimuli provide a strong constraint on the range of the leak rate that matches behavioral data because increased leak rates lead to lower contribution of earlier stimulus fluctuations to choice. We found that, while the fitted leak rate was statistically greater than zero (Fig. 6B; identity task, *t*(11) = 3.01, *p* = 0.012; expression task, *t*(6) = 2.99, *p* = 0.024), it was consistently small across subjects (identity task, mean±s.e.m. across subjects, 0.013 ± 0.004s^−1^; expression task, 0.005 ± 0.002s^−1^). These leak rates correspond to integration time constants larger than 100 s, which is much longer than the duration of each trial (~1 s), supporting near-perfect integration over time.

The second extension allows time-varying sensitivity to sensory evidence within a trial (Levi et al. 2018), as opposed to the constant sensitivity assumed in our main model. To capture a wide variety of plausible temporal dynamics, we added linear and quadratic temporal modulations of drift rate over time to the model (Fig. 6C; Eq. 10). However, the modulation parameters were quite close to zero (Fig. 6D; identity task: *γ*_1_ = −0.16 ± 0.084, *t*(11) = −1.88, *p* = 0.087, *γ*_2_ = 0.027±0.032, *t*(11) = 0.85, *p* = 0.41; expression task: *γ*_1_ = −0.15±0.17, *t*(6) = −0.86, *p* = 0.43, γ_2_ = 0.029 ± 0.085, *t*(6) = 0.34, *p* = 0.75, two-tailed t-test), suggesting a lack of substantial temporal dynamics. Note, however, that the models with slow modulations of sensitivity can not capture the very fast modulations in the observed psychophysical kernels. Although the fits might improve by allowing fast modulations of sensitivity (5-10Hz), we are unaware of sensory or decision mechanisms that can create such fast fluctuations. These fluctuations likely arise from noise due to finite samples in our dataset. Overall, the temporal properties of the decision-making process are consistent with linear multi-feature integration with largely static sensitivities.

### Testing the sequence of spatiotemporal integration

In the models above, temporal integration operates on the momentary evidence generated from the spatial integration of features of each stimulus frame. But is it necessary that spatial integration precedes temporal integration? Although our data-driven analyses above suggest that subjects combined information across facial features (Figs. 3 and 4), it might be plausible that spatial integration follows the temporal integration process instead of preceding it. Specifically, the evidence conferred by each informative facial feature may be independently integrated over time, and then a decision may be rendered based on the collective outcome of the three feature-wise integration processes (i.e., spatial integration following temporal integration). A variety of spatial pooling rules may be employed in such a model. A choice can be determined by the first feature integrator that reaches its bound (Fig. 7A), by the majority of feature integrators, or by the weighted sum of the decision variables of the integrators after the first bound crossing (Fig. 7-1). In all of these model variants, the choice is shaped by multiple features due to the stochasticity of the stimuli and noise (Otto and Mamassian 2012). For example, the “eyes” integrator would dictate the choice in many trials of the identity categorization task, but the other feature integrators would also have a smaller but tangible effect, compatible with the differential contribution of features to choice, as shown in the previous sections (Fig. 2E-F). Are such parallel integration models compatible with the empirical data?

**Figure 7:**
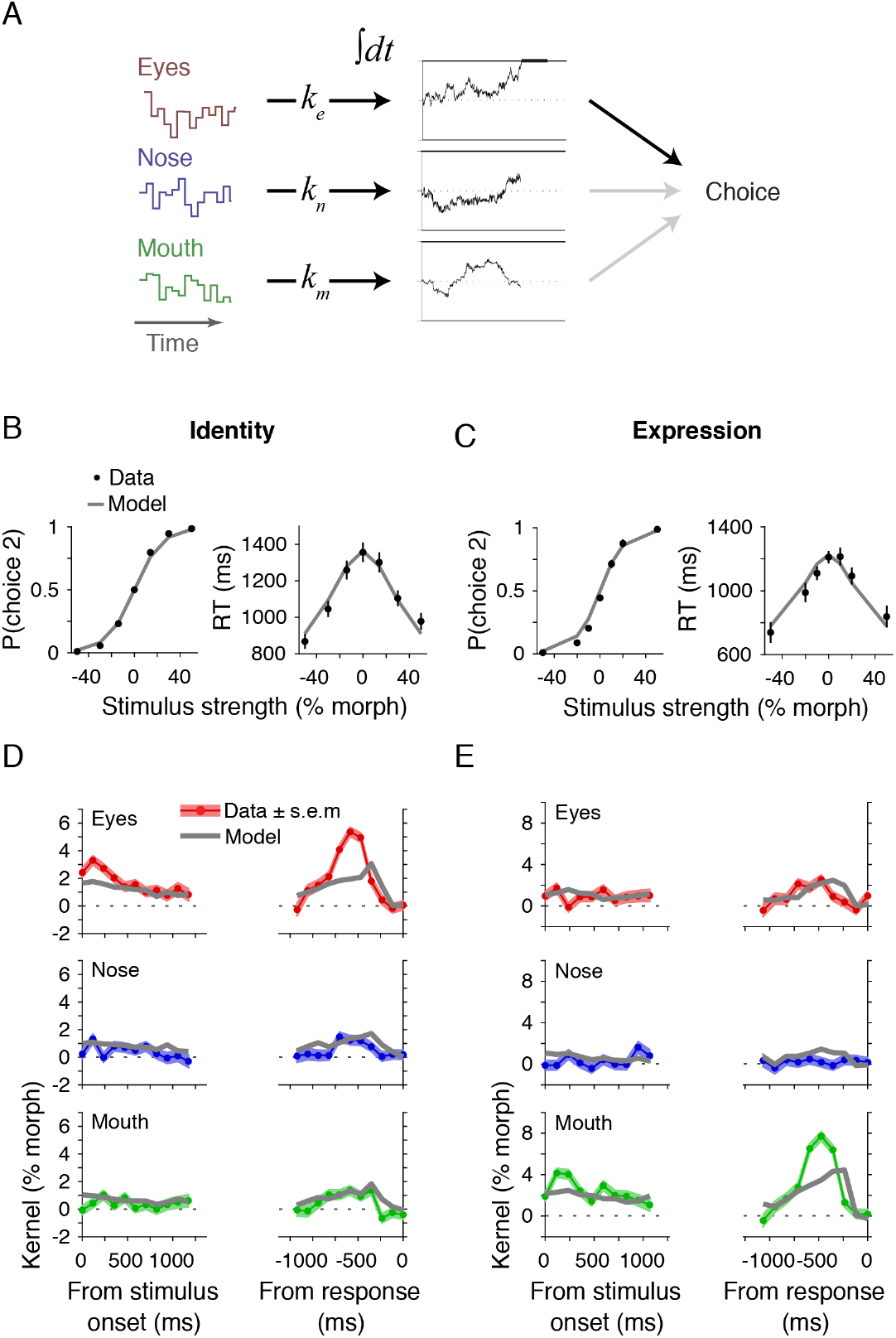
Models with late spatial integration across features fail to explain the experimental data. (**A**) Schematic of a parallel integration model in which single features are independently integrated over time to a decision bound, followed by spatial integration across features to determine the choice. A decision is made based on the first integrator that reaches the decision bound. Models with different decision rules —such as the choice supported by the majority of feature integrators or a linear combination of decision variables across integrators— are examined in Fig. 7-1. (**B, C**) Model fits to psychometric and chronometric functions. Conventions are the same as in Fig. 5B, C. (**D, E**) The model fails to explain psychophysical kernels. Notable discrepancies with the data are visible in the kernels for eyes in the identity task and the kernels for mouth in the expression task.

Figure 7 demonstrates that models with late spatial integration fail to explain the behavior. Although these models could fit the psychometric and chronometric functions (Fig. 7B, C), they underperformed our main model (model log-likelihood difference for the joint distribution of choice and RT, –475.5 in the identity task and –307.6 in the expression task). Moreover, and critically, they did not replicate the subjects’ psychophysical kernels (Fig. 7D, E; identity task: *R*^2^ = 0.46, expression task: *R*^2^ = 0.51). They systematically underestimated saccade-aligned kernel amplitudes for the dominant feature of each task (eyes for identity categorization, and mouth for expression categorization). Further, the predicted model kernels peaked closer to the saccade onset than the empirical kernels. Since psychophysical kernel amplitude is inversely proportional to the decision bound (Okazawa et al. 2018), the lower amplitude of these model kernels suggests that the model overestimated the decision bound, which necessitated a shorter non-decision time to compensate for the elongated decision times caused by the higher bounds. These shorter non-decision times pushed model kernel peaks closer to the saccade onset.

In general, late spatial integration causes a lower signal to noise ratio and is therefore more prone to wrong choices because it ignores part of the available sensory information by terminating the decision-making process based on only one feature or by sub-optimally pooling across spatial features after the termination. To match subjects’ high performance, these models would therefore have to alter the speed-accuracy tradeoff by pushing their decision bound higher than those used by the subjects. However, this change leads to qualitative distortions in the psychophysical kernels. Our approach to augment standard choice and RT fits with psychophysical kernel analyses was key to identify these qualitative differences (Okazawa et al. 2018), which can be used to reliably distinguish models with different orders of spatial and temporal integration.

### What underlies differential contribution of facial features to choice: visual discriminability or decision weight?

The psychophysical kernels and decision-making models in the previous sections indicated that subjects’ choices were deferentially sensitive to fluctuations of the three informative features in each categorization task (Figs. 2E-F, 3C-D, 4, and 8A) and across tasks (*F*(2,51) = 47.4, *p* = 2.3×10^−12^, two-way ANOVA interaction between features and tasks). However, as explained earlier, a higher overall sensitivity to a feature could arise from better visual discriminability of changes in the feature or a higher weight applied by the decision-making process to the feature (Fig. 8B). Both factors are likely present in our task. Task-dependent changes of feature sensitivities support the existence of flexible decision weights. Differential visual discriminability is a likely contributor too because of distinct facial features across faces in the identity task or expressions in the expression task. To determine the exact contribution of visual discriminability and decision weights to the overall sensitivity, we measured the discrimination performance of the same subjects for each facial feature using two tasks: odd-one-out discrimination (Fig. 8C) and categorization of single facial features (Fig. 8-1).

**Figure 8:**
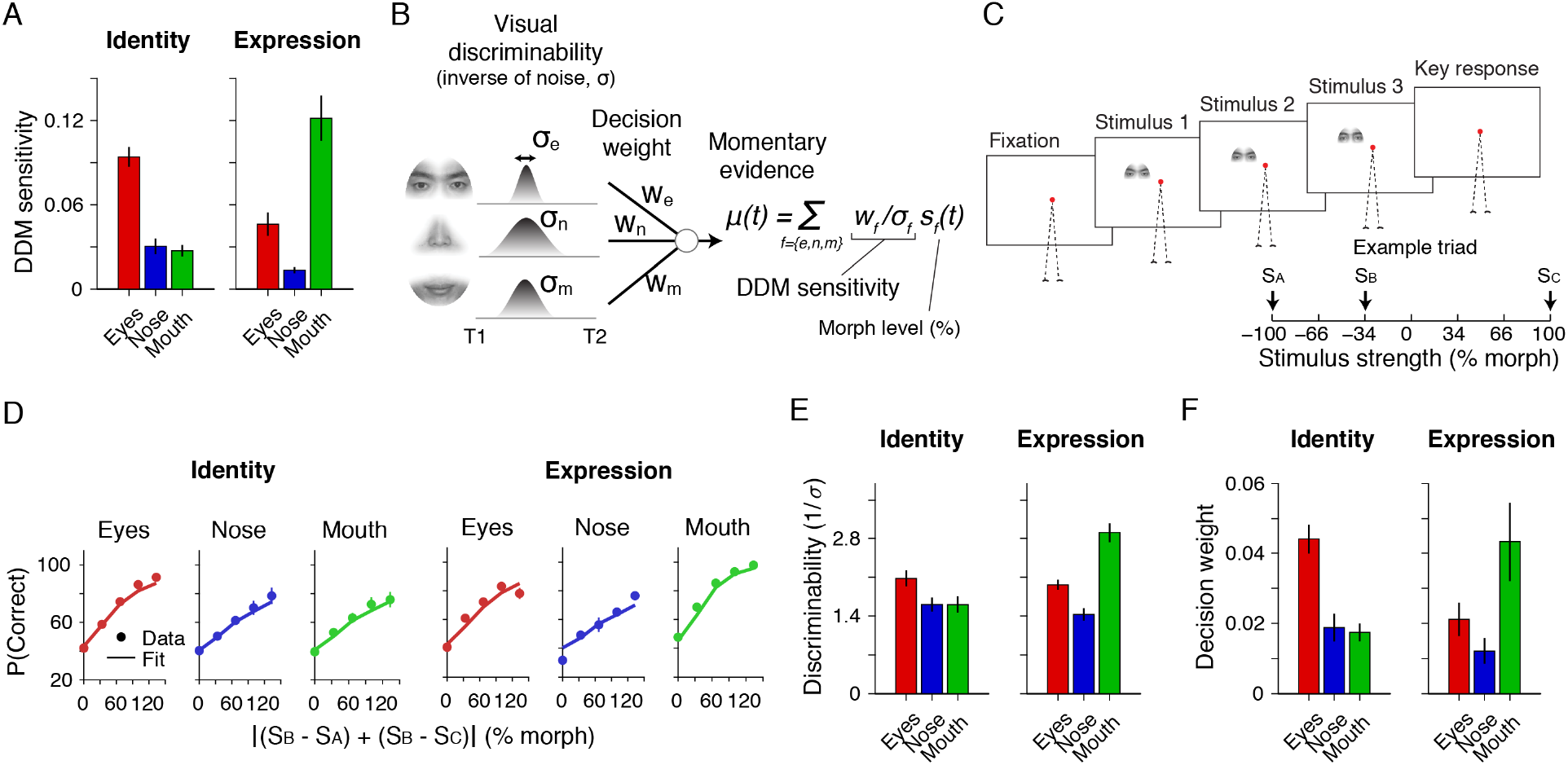
Differential sensitivity of decisions to facial features arises from a combination of visual discriminability and decision weights. (**A**) The sensitivity parameters of the multi-feature drift diffusion model (Fig. 5A) for the informative features in the identity (left) and expression (right) categorization tasks. (**B**) Schematic of the factors that shape differential sensitivities to the informative features. Feature sensitivity in a task arises from two factors: 1) visual discriminability of different morph levels of the feature, and 2) the weight that the decision-making process attributes to the feature. These factors are distinct as visual discriminability arises from the precision of sensory representations, whereas decision weights are flexibly set to achieve a particular behavioral goal. We define visual discriminability of a feature as the inverse of representational noise in units of %morph (*σ_f_*, where *f* could be *e, n*, or *m* for eyes, nose, and mouth, respectively), and the overall sensitivity to a feature as its visual discriminability (1/*σ_f_*) multiplied by the decision weight (*w_f_*). To determine the relative contribution of these two factors, one needs to measure the visual discriminability of facial features. (**C**) The design of an odd-one-out discrimination task to measure the visual discriminability of individual features. In each trial, subjects viewed a sequence of three images of the same feature and reported the one the was perceptually most distinct from the other two. The three images had distinct morph levels (*S_A_, S_B_, S_C_* sorted in ascending order). All subjects of our main task participated in this experiment. (**D**) Subjects’ accuracy as a function of the “distinctness” of the correct stimulus from the other two. Distinctness was quantified as |(*S_B_* – *S_A_*) + (*S_B_* – *S_C_*)|. The lines are the fits of an ideal observer model (see Methods). The accuracy for 0% distinctness was slightly larger than a random choice (33%) because subjects were slightly less likely to choose the middle morph level (*S_B_*). (**E**) Visual discriminability (1/*σ*) estimated using the ideal observer model. Facial features have different discriminability with the eyes slightly more discriminable for the faces used in the identity categorization task and the mouth more discriminable in the expression categorization task. However, these differences in discriminability across features are less pronounced than those in the overall sensitivity (A). (**F**) Decision weights (*w* in B) calculated by dividing the overall sensitivity to the visual discriminability of each feature.

In the odd-one-out task, subjects viewed three consecutive images of a facial feature (eyes, nose, or mouth) with different stimulus strengths and chose the one that was perceptually distinct from the other two (Fig. 8C). Subjects successfully identified the morph level that was distinct from the other two and had higher choice accuracy when the morph level differences were larger (Fig. 8D). However, the improvement of accuracy as a function of morph level difference was not identical across features. The rate of increase (slope of psychometric functions) was higher for the eyes of the identity-task stimuli, and higher for the mouth of the expression-task stimuli, suggesting that the most sensitive features in those tasks were most discriminable, too. We used a model based on signal detection theory (Maloney and Yang 2003) to fit the psychometric functions and retrieve the effective representational noise (*σ_f_*) for each facial feature (see Methods). As expected from the psychometric functions (Fig. 8D), visual discriminability, defined as the inverse of the representational noise in the odd-one-out task, was slightly higher for the eyes of the identity-task stimuli, and for the mouth of the expression-task stimuli (Fig. 8E; identity task: *F*(2,16) = 7.4, *p* = 0.0054, expression task: *F*(2, 4) = 22.7, *p* = 0.0066, repeated measures ANOVA; see Fig. 8-2B for individual subjects).

Similar results were also obtained in the single-feature categorization tasks, where subjects performed categorizations similar to the main task while viewing only one facial feature (Fig. 8-1A). We derived the model sensitivity for each facial feature by fitting a drift diffusion model to the subjects’ choices and RTs (Fig. 8-1B). Because subjects discriminated a single feature in this task, differential weighting of features could not play a role in shaping their behavior, and the model sensitivity for each feature was proportional to the feature discriminability. The order of feature discriminability was similar to that from the odd-one-out task, with eyes showing more discriminability for the stimuli of the identity task and mouth showing more discriminability for the stimuli of the expression task (Fig. 8-1C).

Although the results of both tasks support that visual discriminability was non-uniform across facial features, this contrast was less pronounced than that of the model sensitivities in the main task (Figs. 8E-F, 8-1C, dividing the model sensitivities by the discriminability revealed residual differences reflecting non-uniform decision weights across features (Fig. 8F; *F*(2,30) = 6.1, *p* = 0.0059, two-way ANOVA, main effect of features) and between the tasks (*F*(2,30) = 10.9, *p* = 2.8 × 10^−4^, two-way ANOVA, interaction between features and tasks). In other words, context-dependent decision weights play a significant role in the differential contributions of facial features to decisions. Furthermore, these weights suggest that subjects rely more on more informative (less noisy) features. In fact, the decision weights were positively correlated with visual discriminability (Fig. 8-2C; *R* = 0.744, *p* = 2.0 × 10^−7^), akin to an optimal cue integration process (Drugowitsch et al. 2014; Ernst and Banks 2002; Oruc et al. 2003). Together, the decision-making process in face categorization involves context-dependent adjustment of decision weights that improves behavioral performance.

## Discussion

Successful categorization or identification of objects depends on elaborate sensory and decision-making processes that transmit and use sensory information to implement goal-directed behavior. The properties of the decision-making process remain underexplored for object vision. Existing models commonly assume instantaneous decoding mechanisms based on linear readout of population responses of sensory neurons (Chang and Tsao 2017; Hung et al. 2005; Majaj et al. 2015; Rajalingham et al. 2015), but they are unable to account for aspects of behavior that are based on deliberation on temporally extended visual information common in our daily environments. By extending a quantitative framework developed for studying simpler perceptual decision (Gold and Shadlen 2007; O’Connell et al. 2012; Palmer et al. 2005; Ratcliff and Rouder 1998; Waskom et al. 2019), we establish an experimental and modeling approach that quantitatively links sensory inputs and behavioral responses during face categorization. We show that human face categorization constitutes spatiotemporal evidence integration processes. A spatial integration process aggregates stimulus information into momentary evidence, which is then integrated over time by a temporal integration process. The temporal integration is largely linear and, because of its long time constants, has minimal or no loss of information over time. The spatial integration is also linear and accommodates flexible behavior across tasks by adjusting the weights applied to visual features. These weights remain stable over time in our task, providing no indication that the construction of momentary evidence or its informativeness changes with stimulus viewing time.

Our approach bridges past studies on object recognition and perceptual decision making by formulating face recognition as a process that integrates sensory evidence over space and time. Past research on object recognition focused largely on feedforward visual processing and instantaneous readout of the visual representations, leaving a conceptual gap for understanding the temporally extended processes that underlie perception and action planning based on visual object information. Several studies have attempted to fill this gap by employing noisy object stimuli (Heekeren et al. 2004; Heidari-Gorji et al. 2021; Philiastides et al. 2014; Philiastides and Sajda 2006; Ploran et al. 2007) or sequential presentation of object features (Jack et al. 2014; Ploran et al. 2007). However, their stimulus manipulations did not allow a comprehensive exploration of both spatial and temporal processes. They either created a one-dimensional stimulus axis that eroded the possibility to study spatial integration across features or created temporal sequences that eroded the possibility to study temporal integration jointly with spatial integration. Our success hinges on a novel stimulus design: independent parametric morphing of individual facial features and subliminal spatiotemporal feature fluctuations within trials. Independent feature fluctuations were key to characterize the integration processes, and the subliminal sensory fluctuations ensured that our stimulus manipulations did not alter subjects’ decision strategy, addressing a fundamental challenge (Murray and Gold 2004) posed to alternative methods (e.g., covering face parts: Gosselin and Schyns 2001; Schyns et al. 2002, but see Gosselin and Schyns 2004).

We used three behavioral measures —choice, reaction time, and psychophysical reverse correlation— to assess the mechanisms underlying the behavior. Some key features of the decision-making process cannot be readily inferred solely from choice and reaction time, e.g., the time constant of the integration process (Ditterich 2006; Stine et al. 2020). However, the inclusion of psychophysical kernels provides a more powerful three-pronged approach (Okazawa et al. 2018) that enabled us to establish differential sensitivities for informative features (Fig. 2E-F), linearity of spatial integration (Fig. 3), long time constants (minimum information loss) for temporal integration (Fig. 6B), static feature sensitivities (Fig. 6D), and failure of late spatial integration in the parallel feature integration models (Fig. 7). The precise agreement of psychophysical kernels between model and data (Fig. 5D-E) reinforces our conclusion that face categorization arises from linear spatiotemporal integration of visual evidence.

Face perception is often construed as a “holistic” process because breaking the configuration of face images —e.g., removing parts (Tanaka and Farah 1993), shifting parts (Young et al. 1987), or inverting images (Yin 1969)— reduces performance for face discrimination (Taubert et al. 2011), categorization (Young et al. 1987), or recognition (Tanaka and Farah 1993). However, the mechanistic underpinnings of these phenomena remain elusive (Richler et al. 2012). The linear spatial integration mechanism has the potential to provide mechanistic explanations for some of these holistic effects. For example, changes in the configuration of facial features could reduce visual discriminability of facial features (Murphy and Cook 2017), disrupt spatial integration (Gold et al. 2012; Witthoft et al. 2016), or cause suboptimal weighting of informative features (Sekuler et al. 2004). Holistic effects can also manifest as impairment in facial part recognition when placed together with other uninformative facial parts (composite face effect; Young et al. 1987). This might arise because face stimuli automatically trigger spatial integration that combines information from irrelevant parts. Our approach offers a quantitative path to test these possibilities using a unified modeling framework — a fruitful direction to pursue in the future.

The linearity of spatial integration over facial features has been a source of controversy in the past (Gold 2014; Gold et al. 2012; Shen and Palmeri 2015). The controversy partly stems from the ambiguity in what visual information contributes to face recognition. Some suggest that local shape information of facial parts accounts for holistic face processing (McKone and Yovel 2009), while others suggest that configural information, such as distances between facial features, gives rise to non-linearities (Shen and Palmeri 2015) and holistic properties (Le Grand et al. 2001; Maurer et al. 2002). Our study does not directly address this question because feature locations in our stimuli were kept largely constant to facilitate morphing between faces. However, our approach can be generalized to include configural information and systematically tease apart spatial integration over feature contents from integration over the relative configuration of features. An ideal decision-making process would treat configural information similar to content information — by linearly integrating independent pieces of information. Although our current results strongly suggest linear integration over feature contents, we remain open to emergent non-linearities for configural information.

Another key finding in our experiments is flexible, task-dependent decision weights for informative features (Fig. 8). Past studies demonstrated preferential use of more informative features over others during face and object categorization (De Baene et al. 2008; Schyns et al. 2002; Sigala and Logothetis 2002). But it was not entirely clear whether and by how much subjects’ enhanced sensitivity stemmed from visual discriminability of features or decision weights. We have shown that the differential model sensitivity for facial features in our tasks could not be fully explained by inhomogeneity of visual discriminability across features, thus confirming flexible decision weights for facial features. Importantly, the weights were proportional to the visual discriminability of features in each task (Fig. 8-2C), consistent with the idea of optimal cue integration that explains multi-sensory integration behavior (Drugowitsch et al. 2014; Ernst and Banks 2002; Oruc et al. 2003). Our observation suggests that face recognition is compatible with Bayesian computations in cue combination paradigms (Fetsch et al. 2013; Gold et al. 2012). It is an important future direction to test whether the recognition of other object categories also conforms to such optimal computations (Kersten et al. 2004). Moreover, neural responses to object stimuli can dynamically change due to adaptation or expectation (Kaliukhovich et al. 2013), which can alter both the sensory and decision-making processes (Mather and Sharman 2015; Witthoft et al. 2018). How decision-making processes adapt to dynamic inputs is another important direction to be explored in the future.

The quantitative characterization of behavior is pivotal in linking computational mechanisms and neural activity. Our results raise questions about where and how the spatiotemporal integration of sensory evidence is implemented in the brain for object vision. Traditionally, such processes have been investigated in sensorimotor and association brain regions (Cisek and Kalaska 2005; Gold and Shadlen 2007; Ratcliff et al. 2003; Schall 2019). Also, the inferotemporal cortex (IT), where the face patch system is located (Hesse and Tsao 2020), has been investigated as a dominantly sensory region (DiCarlo et al. 2012; Tanaka 1996). Such a functional division would suggest that IT provides the input (feature information) to the integrators implemented by the downstream frontoparietal circuits. Although feature information is encoded in the frontoparietal circuits (Okazawa et al. 2021), IT neurons also encode combinations of facial features (Chang and Tsao 2017; Freiwald and Tsao 2010) and can flexibly alter their sensitivity based on task demands (Koida and Komatsu 2007), providing a basis for spatial integration. Further, IT neurons represent upcoming choices and show response dynamics that can reflect temporally extended decisions (Akrami et al. 2009; Tajima et al. 2017). An appealing hypothesis is that IT exceeds a purely sensory role and object recognition behaviors arise from interactions between the IT cortex and downstream sensorimotor and association areas. Our experimental framework provides a foundation for studying such interactions by determining the properties of spatiotemporal integration and making quantitative predictions about the underlying neural responses.

## Acknowledgements

The authors would like to thank Stanley J. Komban, Michael L. Waskom, and Koosha Khalvati for helpful discussions. This work was supported by the Simons Collaboration on the Global Brain (grant 542997), McKnight Scholar Award, Pew Scholarship in the Biomedical Sciences, and National Institute of Mental Health (R01 MH109180-01). G.O. was supported by post-doctoral fellowships from the Charles H. Revson Foundation and the Japan Society for the Promotion of Science.

## Extended Data

**Figure 1-1:**
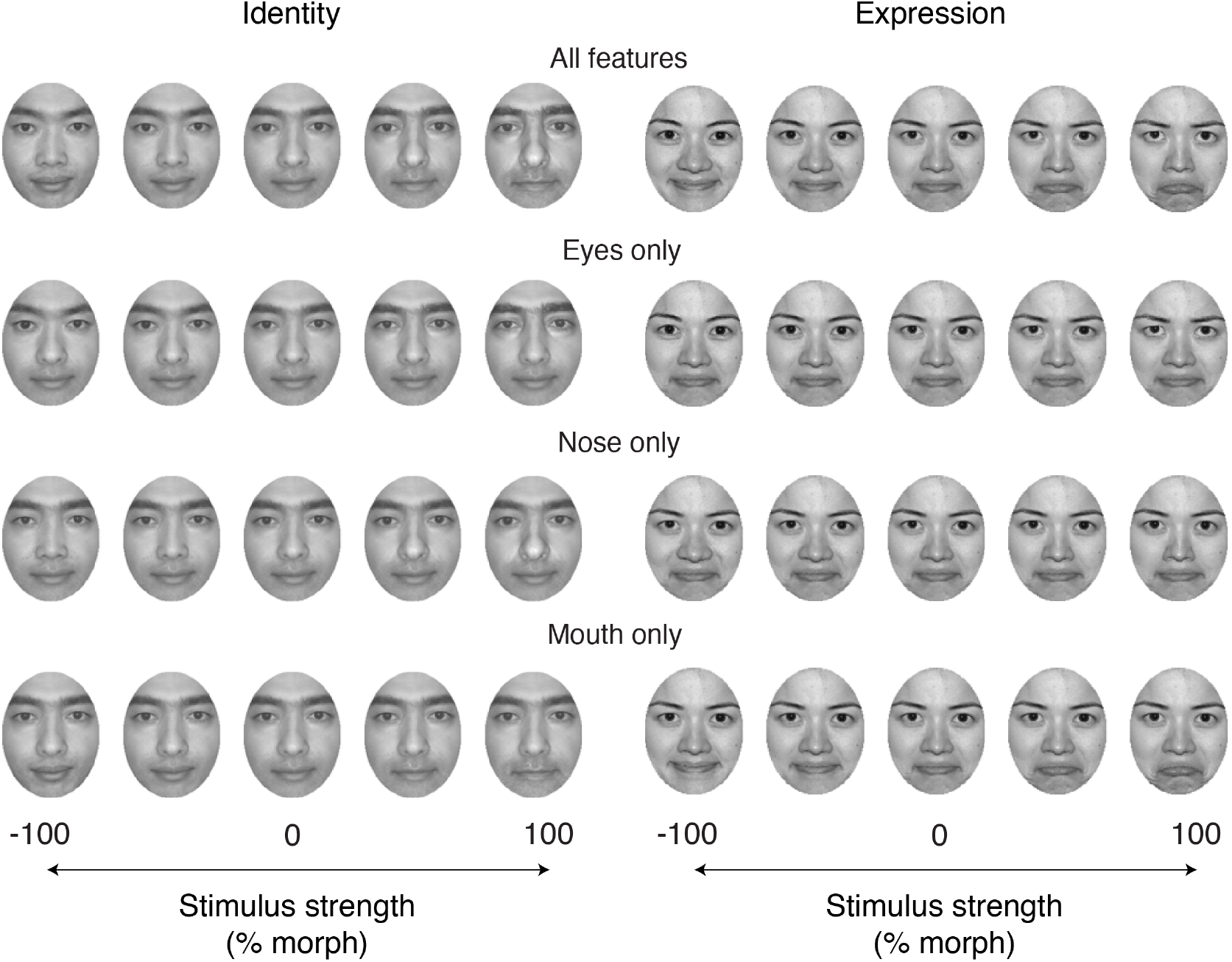
Example face images created using our morph algorithm. Our algorithm independently morphs three facial features (eyes, nose, mouth) between two prototype images. The top row shows a morph continuum where all facial features vary together between two prototypes. Each of the bottom three rows shows a morph continuum for one facial feature, keeping the rest of features constant at 0% morph level. The face size was approximately 83 (W) × 108 (H) pixels (2.18° × 2.83°on the screen). The sizes of individual features were approximately 80 × 28 pixels (eyes), 36 × 30 pixels (nose), and 67 × 22 pixels (mouth). The morphing happened in those local regions, but as can be seen, the resulting images create a seamless continuum of naturalistic faces without noticeable aliasing. The prototype faces in the experiments were chosen from the photographs of MacBrain Face Stimulus Set (Tottenham et al. 2009). For the identity stimuli shown in the figure, we used faces of two authors to avoid copyright issues. For the expression stimuli, the use of the images has been permitted.

**Figure 2-1:**
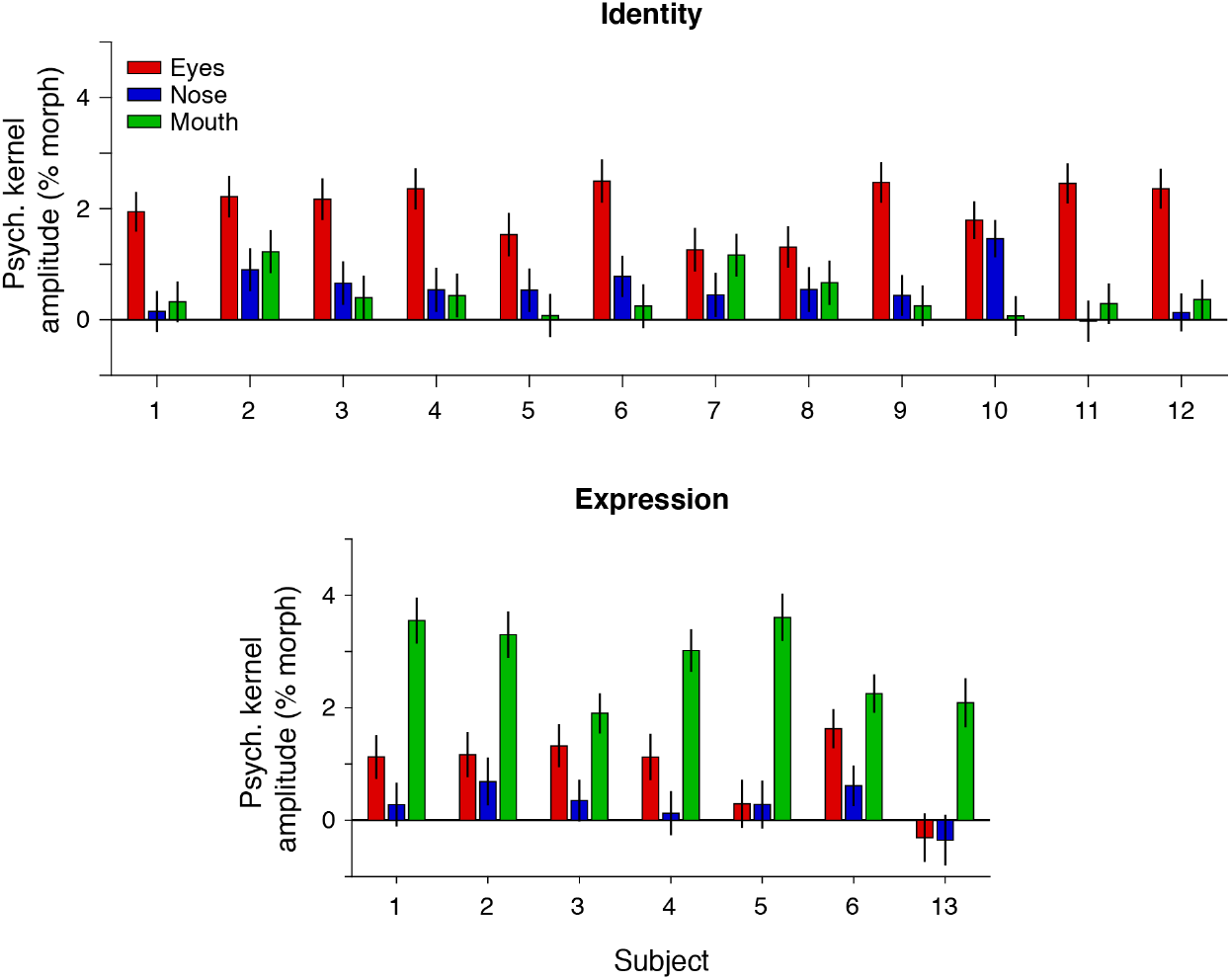
The amplitudes of psychophysical kernels of individual subjects. Individual subjects’ plots corresponding to Fig. 2E-F. Most subjects showed positive kernels for multiple facial features in both identity and expression tasks, indicating that they used multiple facial features for decisions. Error bars indicate s.e.m. across trials. The same subject IDs are consistently used throughout the paper (Figs. 2-1, 3-1, 5-1, 5-2, 8-2).

**Figure 3-1:**
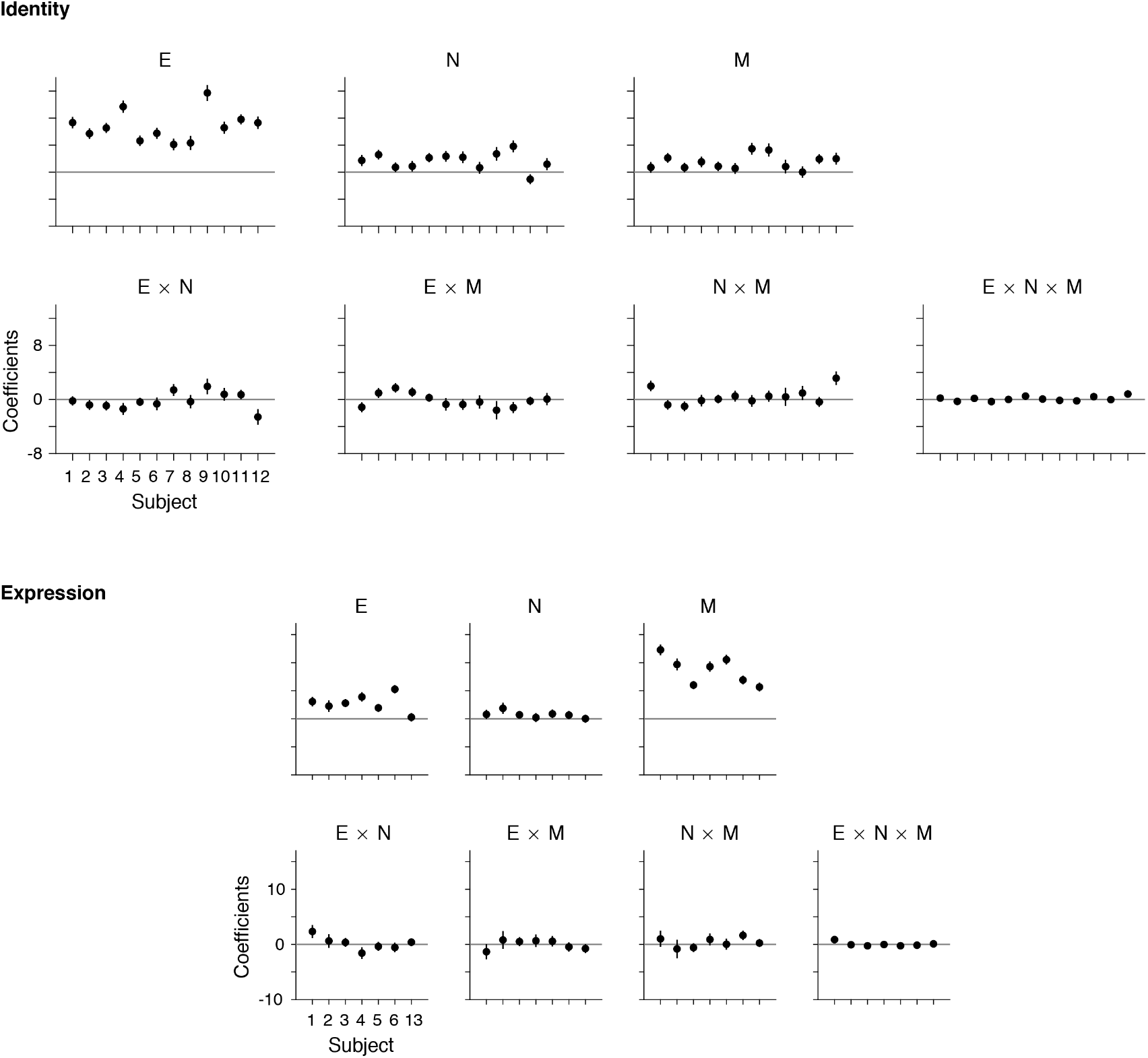
Linear spatial integration of features in individual subjects. Individual subjects’ plots corresponding to Fig. 3C, D. The coefficients for multiplicative factors are close to zero in most subjects, indicating that they integrated spatial evidence in a largely linear fashion. E, N, M indicates eyes, nose, mouth, respectively. Error bars are s.e.m. across trials.

**Figure 5-1:**
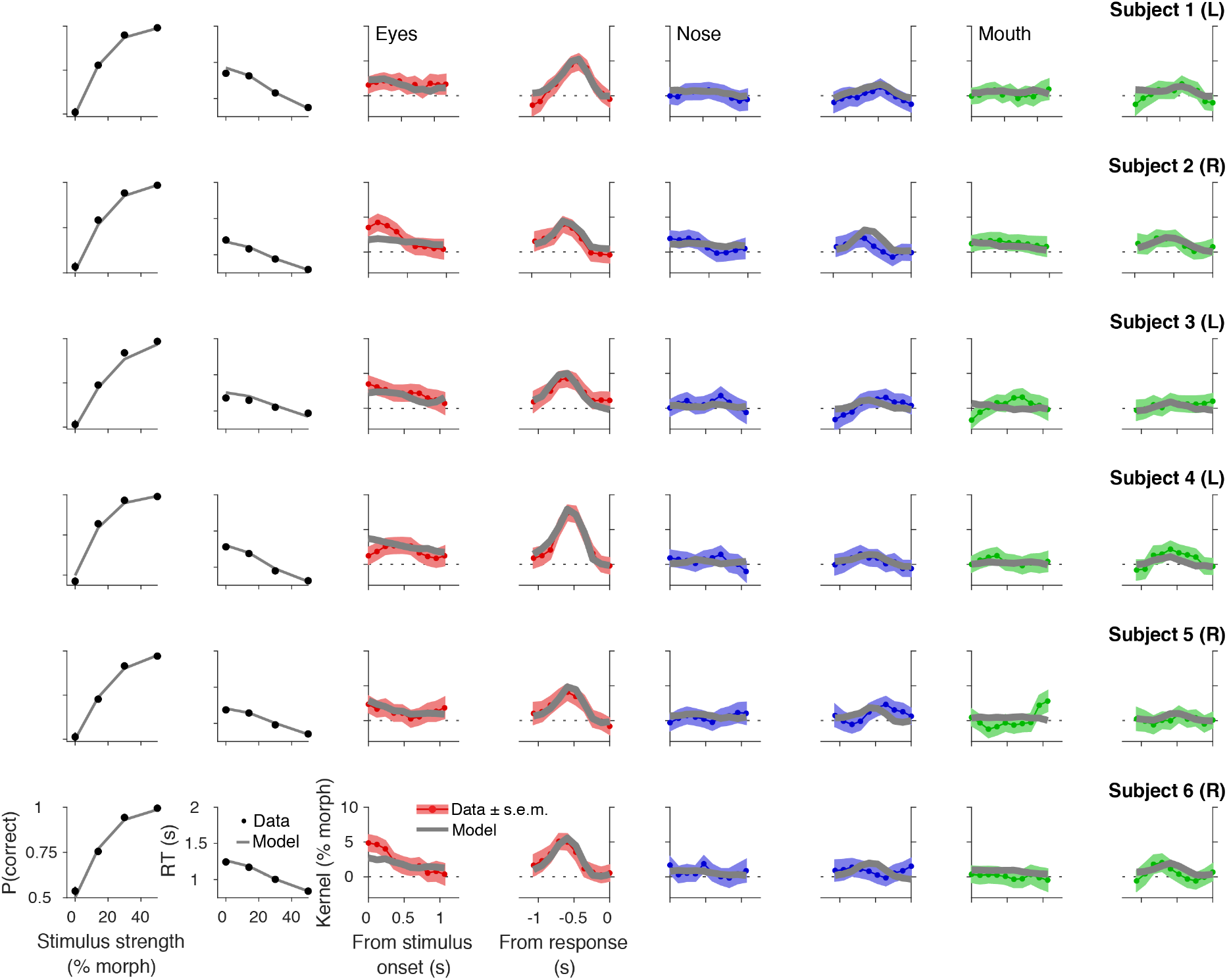
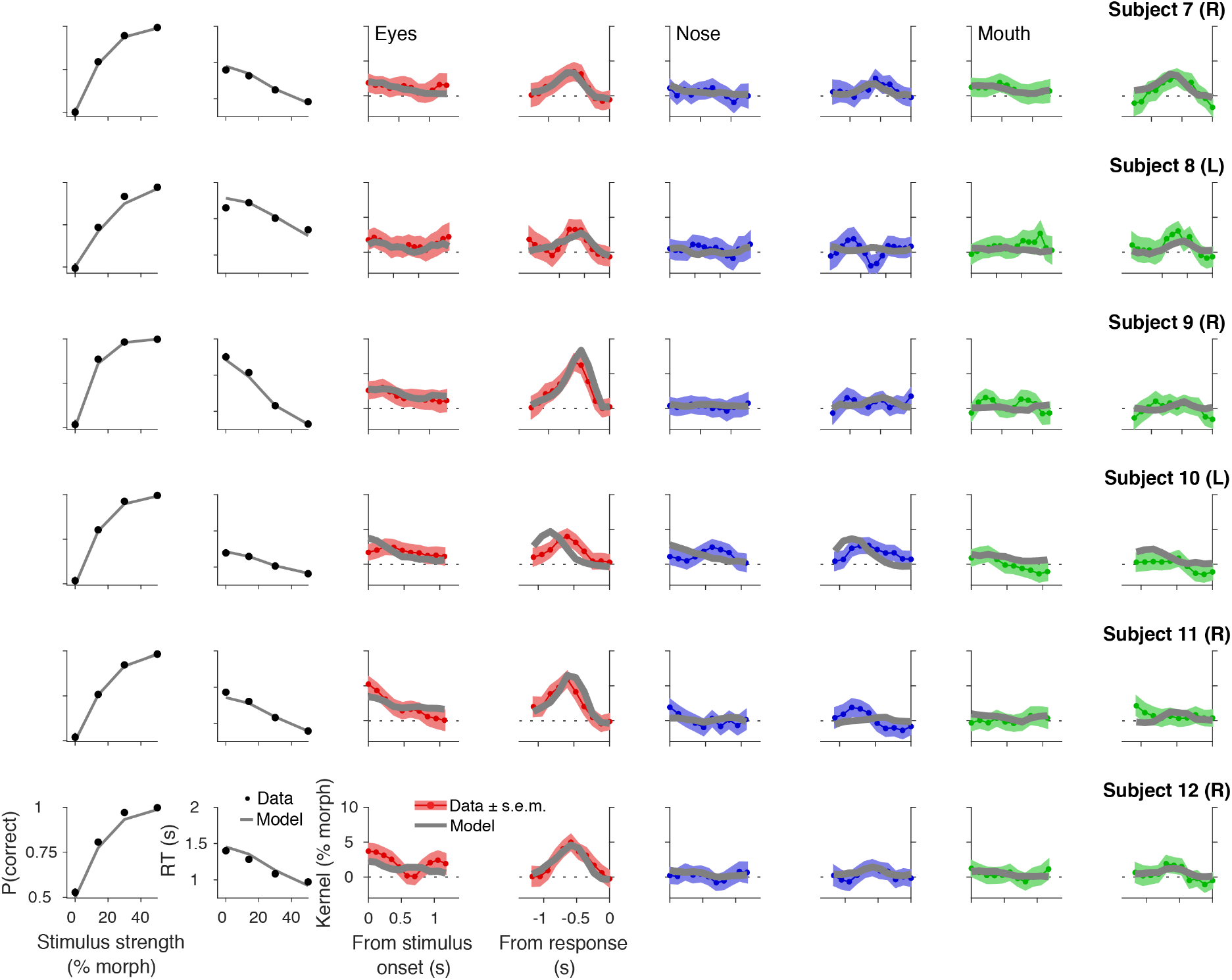
Choice, reaction time, and psychophysical kernels for individual subjects in the identity task. Each row corresponds to the results of one subject. The conventions are the same as in Fig. 5B-E. The multi-feature drift diffusion model with linear spatiotemporal evidence integration (Fig. 5A) fits quite well to behavioral data for most subjects (gray lines). Error bars (s.e.m. across trials) in psychometric and chronometric functions are smaller than the data points. The labels (L or R) next to subjects’ IDs indicate the side on which the parafoveal stimuli were presented for each subject (left or right of the fixation point; see Methods). There was no systematic difference in behavior between subjects who were tested with stimuli on the opposite sides. For the subjects who performed both the identity and expression tasks, the stimuli were always presented on the same side in both tasks. This removes the possibility that stimulus position shapes task-dependent feature weighting. The figure continues to the next page.

**Figure 5-2:**
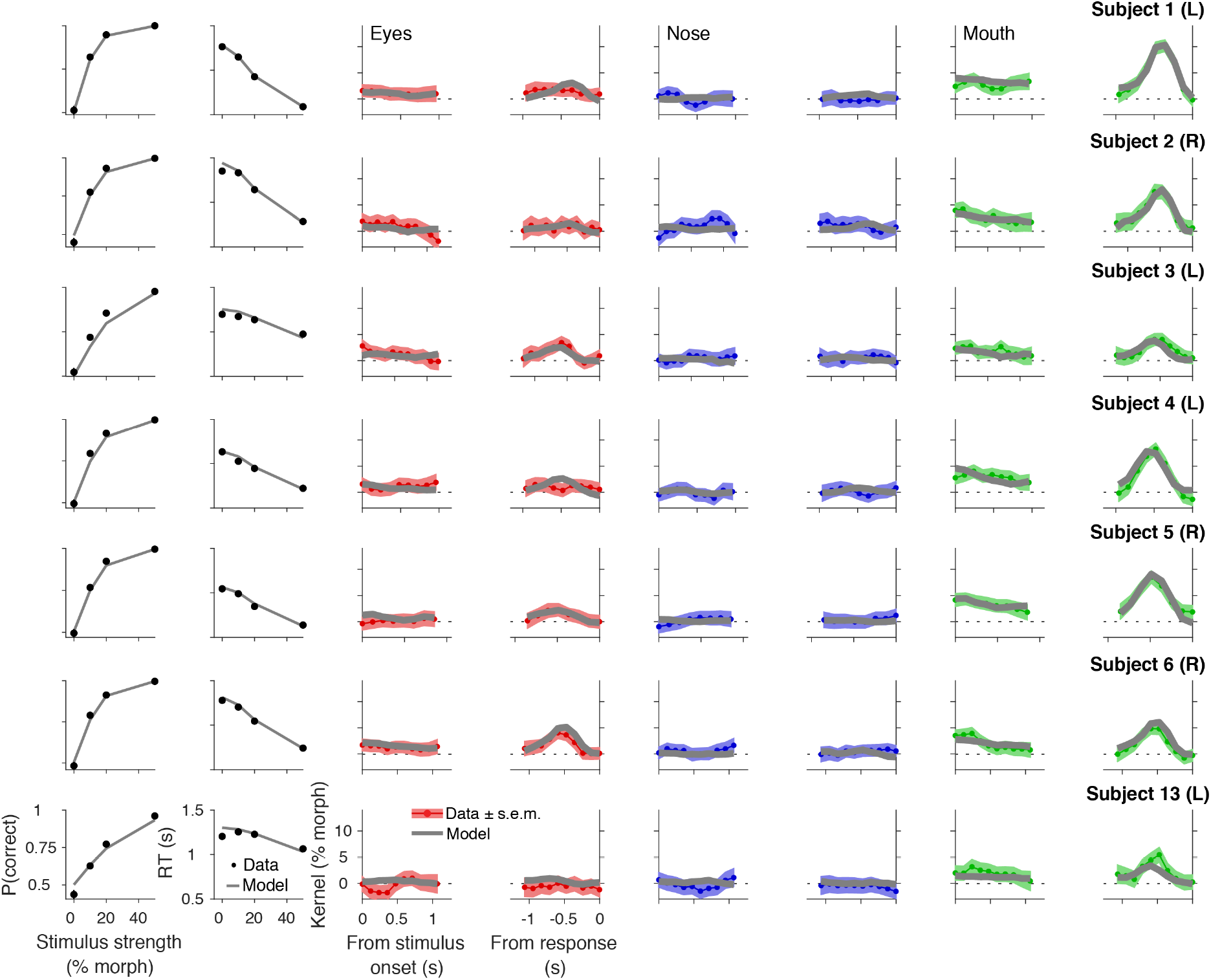
Choice, reaction time, and psychophysical kernels for individual subjects in the expression task.

**Figure 5-3:**
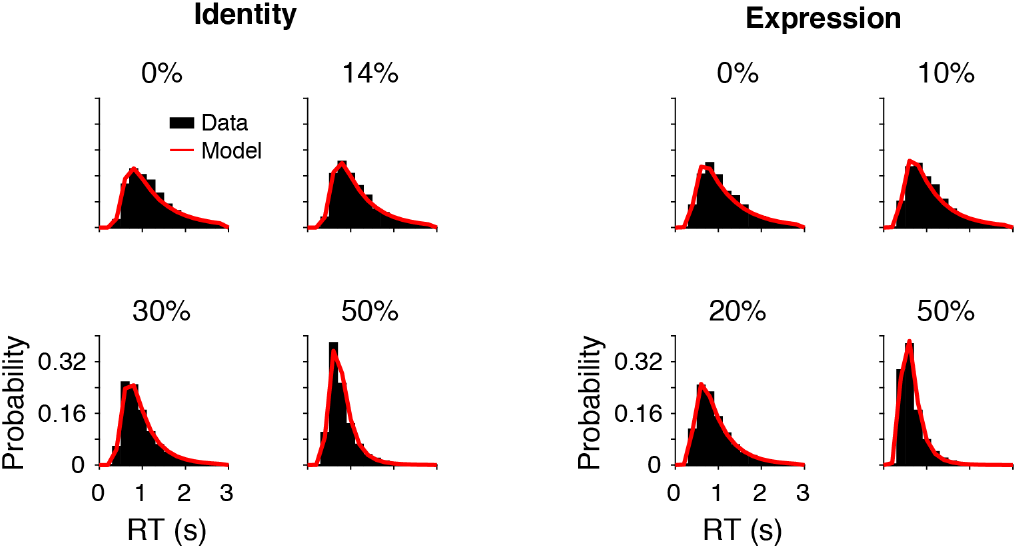
Comparison of reaction time distributions between the behavioral data and our main model. Black bars are the reaction time distributions aggregated across subjects for the specified stimulus strength. Red lines are the distribution generated by the multi-feature drift diffusion model (Fig. 5A).

**Figure 7-1:**
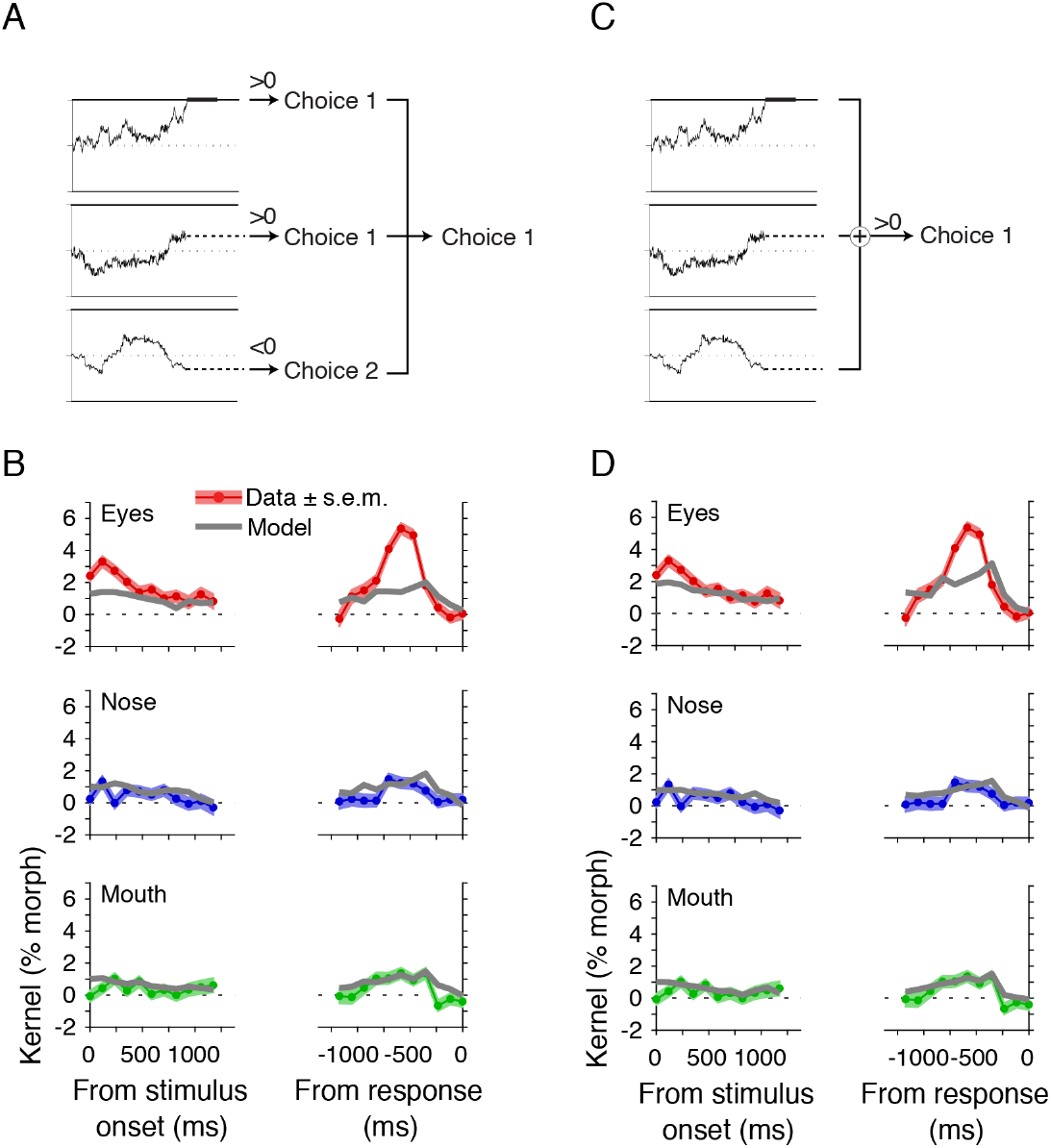
Parallel accumulation models with different decision rules fail to account for behavioral data. (**A**) A parallel integration model that independently accumulates evidence of the three facial features and commits to a choice favored by the majority of the integrators. When one of the three integrators reaches a bound, the model determines the preferred choice of each integrator based on the sign of its decision variable. The model chooses the option supported by the majority of the integrators (i.e., two or more out of three). (**B**) The model in A fails to account for the dynamics of psychophysical kernels. The plots show the results for the identity task. (**C**) Another variant of the parallel integration model that renders a decision based on the summed decision variables of the three integrators. When one integrator reaches a bound, the decision variables of the three accumulators are added, and a decision is made based on the sign of the total evidence. (**D**) The model in C also fails to account for the dynamics of psychophysical kernels in the identity task.

**Figure 8-1:**
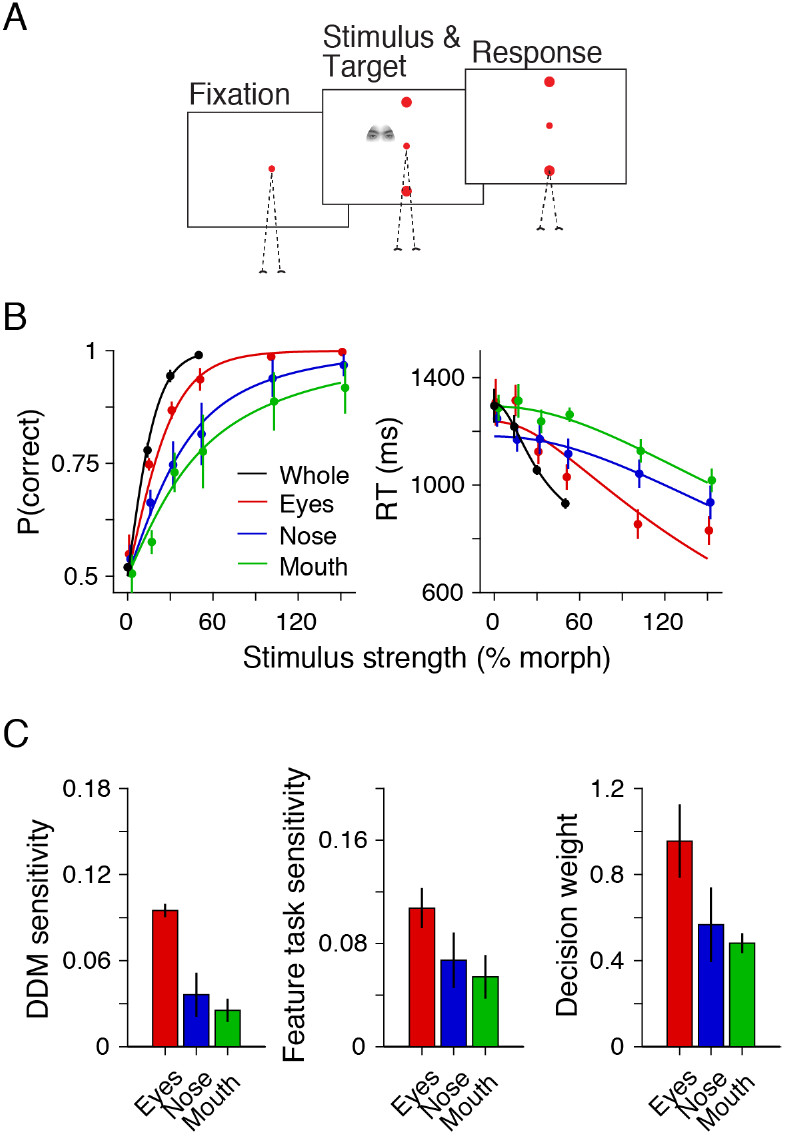
Visual discriminability assessed using a single feature categorization task supports non-uniform decision weights in the main task. (**A**) To quantify visual discriminability for individual facial features, we performed a single feature categorization task with a subset of subjects (4/12 in the identity task). The subjects categorized the facial identities as in the identity task but based their decisions only on one facial feature shown on each trial. The task structure was the same as that of the identity task. Trials of the three facial features were randomly interleaved. To capture the full extent of psychometric functions, we used morph levels ranging from −150% to +150% (see B). The stimuli beyond 100% indicate extrapolation from the prototypes, but the extrapolated images looked natural within the tested range. The task was performed in the same sessions with the main face categorization task (total: 5,571 trials). (**B**) Psychometric and chronometric functions for each facial feature. As a comparison, the same subjects’ performance in the identity task is shown in black. The lines are the fits of a drift diffusion model for each facial feature. (**C**) Comparison of the model feature sensitivities in the main task and in the single feature categorization task. The model feature sensitivities computed from the main identity task (Fig. 8A) can reflect visual discriminability of individual facial features and decision weights dependent on task demands (Fig. 8B), whereas the model sensitivities obtained in the single feature task should reflect only visual discriminability. Dividing the sensitivities of the main task by the single-feature sensitivities recover decision weights, which revealed unequal weighting over features (right). The pattern is consistent with the results based on the discrimination performance in the odd-one-out task (Fig. 8E-F).

**Figure 8-2:**
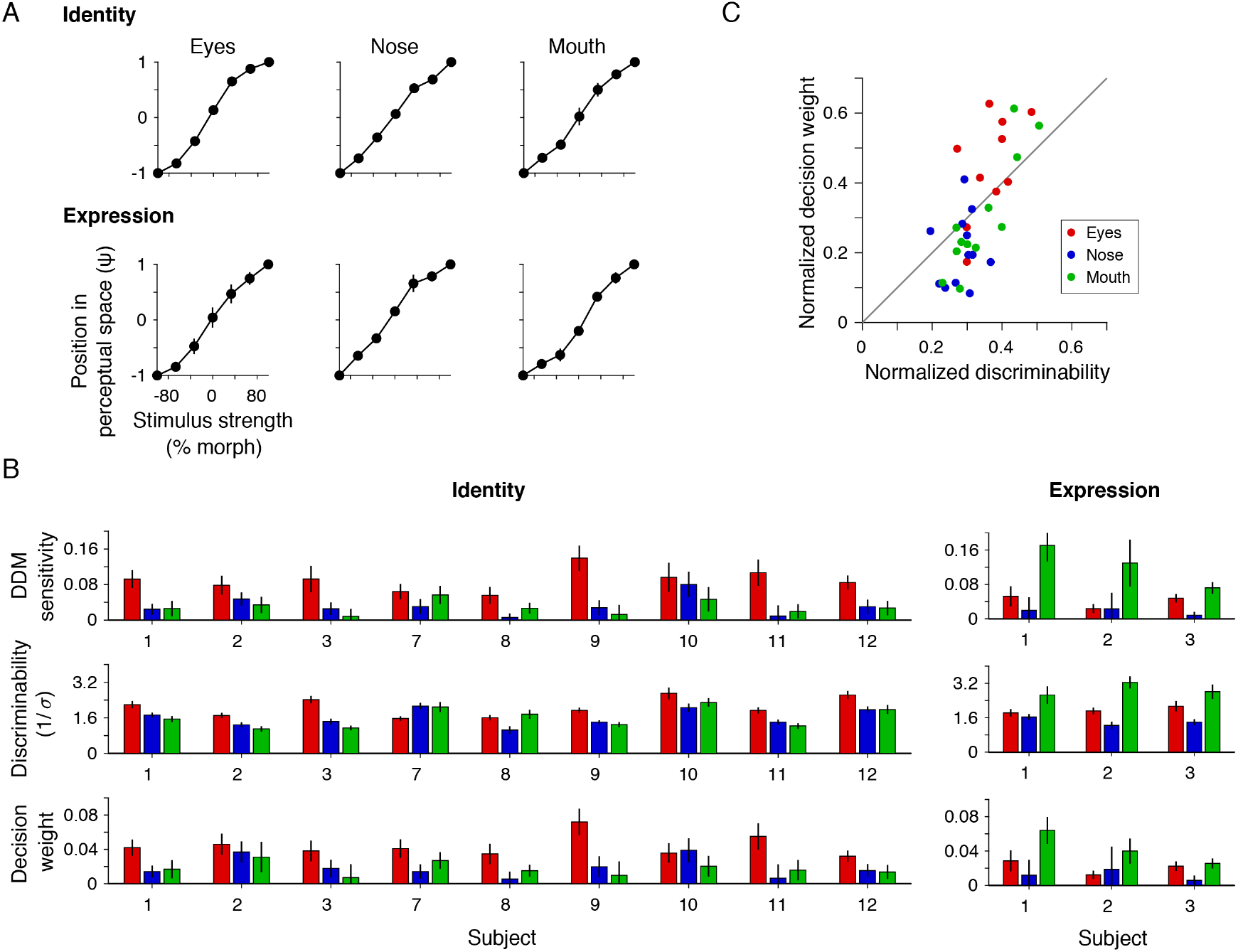
The odd-one-out discrimination task reveals differential visual discriminability for informative features, which were proportional to decision weights. (**A**) The subjects’ stimulus discriminability in the odd-one-out task can be largely explained using the Euclidean distance of morph levels between stimuli. We explained the subjects’ responses in the odd-one-out task (Fig. 8D) based on the representational noise of a feature (σ) and the relative distance between individual pairs of stimuli in a perceptual space (*ψ_i_* and *ψ_j_* for stimuli *i* and *j*). The *ψ* of all intermediate morph levels and *σ* were obtained using a maximum likelihood estimation (see Methods). The *ψ* of the two morph ends were anchored to −1 and +1 to avoid redundant degrees of freedom in the fits. The resulting *ψ* (see the panel) are roughly linearly related to morph levels, indicating that the perceptual distances between stimuli could be largely explained using the Euclidean distance of the stimulus strengths. This linearity ensures that one can use the inverse of the representational noise (1/*σ*) as a metric for the visual discriminability of each facial feature. (**B**) The individual subjects’ plots corresponding to Fig. 8A, E-F. The top panels show the model sensitivities for individual facial features in the main task (*k_e_, k_n_, k_m_* in Fig. 5A). The middle panels show the visual discriminability (1/*σ*) of each feature in the odd-one-out task. Dividing the sensitivity (top) by the discriminability (middle) reveals the contribution of decision weights to the model sensitivities in face categorization (bottom). The results show that most subjects non-uniformly weighted facial features during face categorization. (**C**) Positive correlation between the visual discriminability (middle panels in B) and the decision weights (bottom panels in B) of features is consistent with optimal cue combination. Each dot corresponds to the values of one facial feature of one subject. The plot aggregates data from both the identity and expression tasks. To optimally combine evidence, weights should be proportional to the discriminability of features (Drugowitsch et al. 2014). The plot indeed revealed a significantly positive correlation (*R* = 0.744, *p* = 2.0 × 10^−7^). Both the discriminability and decision weights are normalized within subjects (the sum of all features is fixed to one) to account for the variability of absolute discriminability and weight across subjects.

**Video 1:**
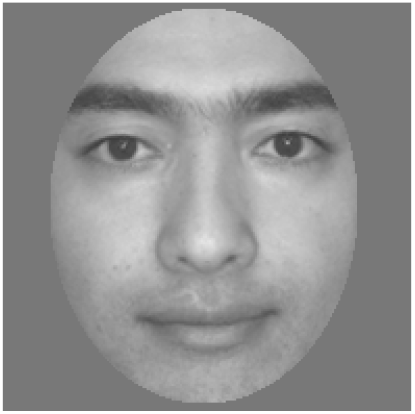
An example dynamic image sequence used in the experiment. The video shows an example image sequence similar to those used in the experiments. The sequence consists of face images interleaved by masks. For each face image, the morph levels of three facial features (eyes, nose, mouth) were randomly sampled from a Gaussian distribution centered on the nominal morph level for the trial (Fig. 1C). The masks made these stimulus fluctuations subliminal. Note that the size and frame rate of the video does not reproduce the actual stimuli used in the experiments.

## References

Ahumada, A. J. (1996). Perceptual classification images from vernier acuity masked by noise. Perception, 25(1-suppl), 2.

Akrami, A., Liu, Y., Treves, A., & Jagadeesh, B. (2009). Converging neuronal activity in inferior temporal cortex during the classification of morphed stimuli. Cereb Cortex, 19(4), 760–76.

Barraclough, N. E., & Perrett, D. I. (2011). From single cells to social perception. Philos Trans R Soc Lond B Biol Sci, 366(1571), 1739–52.

Bogacz, R., Brown, E., Moehlis, J., Holmes, P., & Cohen, J. D. (2006). The physics of optimal decision making: A formal analysis of models of performance in two-alternative forced-choice tasks. Psychological review, 113(4), 700–765.

Brainard, D. H. (1997). The psychophysics toolbox. Spat Vis, 10(4), 433–6.

Brunton, B. W., Botvinick, M. M., & Brody, C. D. (2013). Rats and humans can optimally accumulate evidence for decision-making. Science, 340(6128), 95–8.

Carlson, T. A., Ritchie, J. B., Kriegeskorte, N., Durvasula, S., & Ma, J. (2014). Reaction time for object categorization is predicted by representational distance. J Cogn Neurosci, 26(1), 132–42.

Chang, L., & Tsao, D. Y. (2017). The code for facial identity in the primate brain. Cell, 169(6), 1013–1028 e14.

Cisek, P., & Kalaska, J. F. (2005). Neural correlates of reaching decisions in dorsal premotor cortex: Specification of multiple direction choices and final selection of action. Neuron, 45(5), 801–14.

De Baene, W., Ons, B., Wagemans, J., & Vogels, R. (2008). Effects of category learning on the stimulus selectivity of macaque inferior temporal neurons. Learn Mem, 15(9), 717–27.

DiCarlo, J. J., & Cox, D. D. (2007). Untangling invariant object recognition. Trends Cogn Sci, 11(8), 333–41.

DiCarlo, J. J., Zoccolan, D., & Rust, N. C. (2012). How does the brain solve visual object recognition? Neuron, 73(3), 415–34.

Ditterich, J. (2006). Stochastic models of decisions about motion direction: Behavior and physiology. Neural Netw, 19(8), 981–1012.

Drugowitsch, J., DeAngelis, G. C., Klier, E. M., Angelaki, D. E., & Pouget, A. (2014). Optimal multisensory decisionmaking in a reaction-time task. Elife, 3, e03005.

Drugowitsch, J., Wyart, V., Devauchelle, A. D., & Koechlin, E. (2016). Computational precision of mental inference as critical source of human choice suboptimality. Neuron, 92(6), 1398–1411.

Ernst, M. O., & Banks, M. S. (2002). Humans integrate visual and haptic information in a statistically optimal fashion. Nature, 415(6870), 429–33.

Fetsch, C. R., DeAngelis, G. C., & Angelaki, D. E. (2013). Bridging the gap between theories of sensory cue integration and the physiology of multisensory neurons. Nat Rev Neurosci, 14(6), 429–42.

Freiwald, W. A., & Tsao, D. Y. (2010). Functional compartmentalization and viewpoint generalization within the macaque face-processing system. Science, 330(6005), 845–51.

Gauthier, I., Anderson, A. W., Tarr, M. J., Skudlarski, P., & Gore, J. C. (1997). Levels of categorization in visual recognition studied using functional magnetic resonance imaging. Curr Biol, 7(9), 645–51.

Gauthier, I., Williams, P., Tarr, M. J., & Tanaka, J. (1998). Training ‘greeble’ experts: A framework for studying expert object recognition processes. Vision Res, 38(15-16), 2401–28.

Gold, J. I., & Shadlen, M. N. (2007). The neural basis of decision making. Annu Rev Neurosci, 30, 535–74.

Gold, J. M. (2014). A perceptually completed whole is less than the sum of its parts. Psychol Sci, 25(6), 1206–17.

Gold, J. M., Mundy, P. J., & Tjan, B. S. (2012). The perception of a face is no more than the sum of its parts. Psychol Sci, 23(4), 427–34.

Gosselin, F., & Schyns, P. G. (2001). Bubbles: A technique to reveal the use of information in recognition tasks. Vision Res, 41(17), 2261–71.

Gosselin, F., & Schyns, P. G. (2004). No troubles with bubbles: A reply to Murray and Gold. Vision Res, 44(5), 471–7, discussion 479–82.

Hanks, T., Kiani, R., & Shadlen, M. N. (2014). A neural mechanism of speed-accuracy tradeoff in macaque area LIP. Elife, 3, e02260.

Heekeren, H. R., Marrett, S., Bandettini, P. A., & Ungerleider, L. G. (2004). A general mechanism for perceptual decision-making in the human brain. Nature, 431(7010), 859–62.

Heidari-Gorji, H., Ebrahimpour, R., & Zabbah, S. (2021). A temporal hierarchical feedforward model explains both the time and the accuracy of object recognition. Scientific Reports, 11(1), 5640.

Heitz, R. P., & Schall, J. D. (2012). Neural mechanisms of speed-accuracy tradeoff. Neuron, 76(3), 616–28.

Hesse, J. K., & Tsao, D. Y. (2020). The macaque face patch system: A turtle’s underbelly for the brain. Nat Rev Neurosci, 21(12), 695–716.

Hung, C. P., Kreiman, G., Poggio, T., & DiCarlo, J. J. (2005). Fast readout of object identity from macaque inferior temporal cortex. Science, 310(5749), 863–6.

Jack, R. E., Garrod, O. G., & Schyns, P. G. (2014). Dynamic facial expressions of emotion transmit an evolving hierarchy of signals over time. Curr Biol, 24(2), 187–92.

Kaliukhovich, D. A., De Baene, W., & Vogels, R. (2013). Effect of adaptation on object representation accuracy in macaque inferior temporal cortex. J Cogn Neurosci, 25(5), 777–89.

Kampf, M., Nachson, I., & Babkoff, H. (2002). A serial test of the laterality of familiar face recognition. Brain and Cognition, 50(1), 35–50.

Kanwisher, N., & Yovel, G. (2006). The fusiform face area: A cortical region specialized for the perception of faces. Philos Trans R Soc Lond B Biol Sci, 361(1476), 2109–28.

Karlin, S., & Taylor, H. E. (1981). A second course in stochastic processes. Elsevier.

Kersten, D., Mamassian, P., & Yuille, A. (2004). Object perception as bayesian inference. Annu Rev Psychol, 55, 271–304.

Kiani, R., & Shadlen, M. N. (2009). Representation of confidence associated with a decision by neurons in the parietal cortex. Science, 324(5928), 759–64.

Koida, K., & Komatsu, H. (2007). Effects of task demands on the responses of color-selective neurons in the inferior temporal cortex. Nat Neurosci, 10(1), 108–16.

Kreichman, O., Bonneh, Y. S., & Gilaie-Dotan, S. (2020). Investigating face and house discrimination at foveal to parafoveal locations reveals category-specific characteristics. Sci Rep, 10(1), 8306.

Le Grand, R., Mondloch, C. J., Maurer, D., & Brent, H. P. (2001). Early visual experience and face processing. Nature, 410(6831), 890.

Levi, A. J., Yates, J. L., Huk, A. C., & Katz, L. N. (2018). Strategic and dynamic temporal weighting for perceptual decisions in humans and macaques. eNeuro, 5(5), ENEURO.0169–18.2018.

Levy, I., Hasson, U., Avidan, G., Hendler, T., & Malach, R. (2001). Center-periphery organization of human object areas. Nat Neurosci, 4(5), 533–9.

Link, S. W. (1992). The wave theory of difference and similarity. Psychology Press.

Majaj, N. J., Hong, H., Solomon, E. A., & DiCarlo, J. J. (2015). Simple learned weighted sums of inferior temporal neuronal firing rates accurately predict human core object recognition performance. J Neurosci, 35(39), 13402–18.

Maloney, L. T., & Yang, J. N. (2003). Maximum likelihood difference scaling. J Vis, 3(8), 573–85.

Mather, G., & Sharman, R. J. (2015). Decision-level adaptation in motion perception. R Soc Open Sci, 2(12), 150418.

Maurer, D., Grand, R. L., & Mondloch, C. J. (2002). The many faces of configural processing. Trends Cogn Sci, 6(6), 255–260.

McKone, E., & Yovel, G. (2009). Why does picture-plane inversion sometimes dissociate perception of features and spacing in faces, and sometimes not? toward a new theory of holistic processing. Psychon Bull Rev, 16(5), 778–97.

Murphy, J., & Cook, R. (2017). Revealing the mechanisms of human face perception using dynamic apertures. Cognition, 169, 25–35.

Murray, R. F., & Gold, J. M. (2004). Troubles with bubbles. Vision Res, 44(5>), 461–70.

O’Connell, R. G., Dockree, P. M., & Kelly, S. P. (2012). A supramodal accumulation-to-bound signal that determines perceptual decisions in humans. Nat Neurosci, 15(12), 1729–35.

Okazawa, G., Hatch, C. E., Mancoo, A., Machens, C. K., & Kiani, R. (2021). Representational geometry of perceptual decisions in the monkey parietal cortex. Cell, in press.

Okazawa, G., Sha, L., Purcell, B. A., & Kiani, R. (2018). Psychophysical reverse correlation reflects both sensory and decision-making processes. Nat Commun, 9(1), 3479.

Oruc, I., Maloney, L. T., & Landy, M. S. (2003). Weighted linear cue combination with possibly correlated error. Vision Res, 43(23), 2451–68.

Otto, T. U., & Mamassian, P. (2012). Noise and correlations in parallel perceptual decision making. Curr Biol, 22(15), 1391–6.

Palmer, J., Huk, A. C., & Shadlen, M. N. (2005). The effect of stimulus strength on the speed and accuracy of a perceptual decision. J Vis, 5(5), 376–404.

Perrodin, C., Kayser, C., Abel, T. J., Logothetis, N. K., & Petkov, C. I. (2015). Who is that? brain networks and mechanisms for identifying individuals. Trends Cogn Sci, 19(12), 783–796.

Philiastides, M. G., Heekeren, H. R., & Sajda, P. (2014). Human scalp potentials reflect a mixture of decision-related signals during perceptual choices. J Neurosci, 34(50), 16877–89.

Philiastides, M. G., & Sajda, P. (2006). Temporal characterization of the neural correlates of perceptual decision making in the human brain. Cereb Cortex, 16(4), 509–18.

Ploran, E. J., Nelson, S. M., Velanova, K., Donaldson, D. I., Petersen, S. E., & Wheeler, M. E. (2007). Evidence accumulation and the moment of recognition: Dissociating perceptual recognition processes using fMRI. J Neurosci, 27(44), 11912–24.

Rajalingham, R., Schmidt, K., & DiCarlo, J. J. (2015). Comparison of object recognition behavior in human and monkey. J Neurosci, 35(35), 12127–36.

Ramon, M., Caharel, S., & Rossion, B. (2011). The speed of recognition of personally familiar faces. Perception, 40(4), 437–49.

Ratcliff, R., Cherian, A., & Segraves, M. (2003). A comparison of macaque behavior and superior colliculus neuronal activity to predictions from models of two-choice decisions. J Neurophysiol, 90(3), 1392–407.

Ratcliff, R., & Rouder, J. N. (2000). A diffusion model account of masking in two-choice letter identification. J Exp Psychol Hum Percept Perform, 26(1), 127–40.

Ratcliff, R., & Rouder, J. N. (1998). Modeling response times for two-choice decisions. Psychological Science, 9(5), 347–356.

Richler, J. J., Palmeri, T. J., & Gauthier, I. (2012). Meanings, mechanisms, and measures of holistic processing. Front Psychol, 3, 553.

Riesenhuber, M., & Poggio, T. (1999). Hierarchical models of object recognition in cortex. Nat Neurosci, 2(11), 1019–25.

Rossion, B. (2014). Understanding face perception by means of human electrophysiology. Trends Cogn Sci, 18(6), 310–8.

Schall, J. D. (2019). Accumulators, neurons, and response time. Trends Neurosci, 42(12), 848–860.

Schyns, P. G., Bonnar, L., & Gosselin, F. (2002). Show me the features! Understanding recognition from the use of visual information. Psychol Sci, 13(5), 402–9.

Schyns, P. G., Petro, L. S., & Smith, M. L. (2007). Dynamics of visual information integration in the brain for categorizing facial expressions. Curr Biol, 17(18), 1580–5.

Sekuler, A. B., Gaspar, C. M., Gold, J. M., & Bennett, P. J. (2004). Inversion leads to quantitative, not qualitative, changes in face processing. Curr Biol, 14(5), 391–6.

Shadlen, M. N., Hanks, T. D., Churchland, A. K., Kiani, R., & Yang, T. (2006). The speed and accuracy of a simple perceptual decision: A mathematical primer. Bayesian brain: Probabilistic approaches to neural coding (pp. 209–37). The MIT Press.

Shen, J., & Palmeri, T. J. (2015). The perception of a face can be greater than the sum of its parts. Psychon Bull Rev, 22(3), 710–6.

Sigala, N., & Logothetis, N. K. (2002). Visual categorization shapes feature selectivity in the primate temporal cortex. Nature, 415(6869), 318–20.

Smith, P. L., & Little, D. R. (2018). Small is beautiful: In defense of the small-n design. Psychon Bull Rev, 25(6), 2083–2101.

Smith, P. L., & Vickers, D. (1988). The accumulator model of two-choice discrimination. Journal of Mathematical Psychology, 32(2), 135–168.

Stine, G. M., Zylberberg, A., Ditterich, J., & Shadlen, M. N. (2020). Differentiating between integration and non-integration strategies in perceptual decision making. Elife, 9, e55365.

Tajima, S., Koida, K., Tajima, C. I., Suzuki, H., Aihara, K., & Komatsu, H. (2017). Task-dependent recurrent dynamics in visual cortex. Elife, 6, e26868.

Tanaka, J. W., & Farah, M. J. (1993). Parts and wholes in face recognition. Q J Exp Psychol A, 46(2), 225–45.

Tanaka, K. (1996). Inferotemporal cortex and object vision. Annu Rev Neurosci, 19, 109–39.

Taubert, J., Apthorp, D., Aagten-Murphy, D., & Alais, D. (2011). The role of holistic processing in face perception: Evidence from the face inversion effect. Vision Res, 51(11), 1273–8.

Thorpe, S., Fize, D., & Marlot, C. (1996). Speed of processing in the human visual system. Nature, 381(6582), 520–2.

Tottenham, N., Tanaka, J. W., Leon, A. C., McCarry, T., Nurse, M., Hare, T. A., Marcus, D. J., Westerlund, A., Casey, B. J., & Nelson, C. (2009). The nimstim set of facial expressions: Judgments from untrained research participants. Psychiatry Res, 168(3), 242–9.

Tsao, D. Y., & Livingstone, M. S. (2008). Mechanisms of face perception. Annu Rev Neurosci, 31, 411–37.

Usher, M., & McClelland, J. L. (2001). The time course of perceptual choice: The leaky, competing accumulator model. Psychological review, 108(3), 550–592.

Waskom, M. L., & Kiani, R. (2018). Decision making through integration of sensory evidence at prolonged timescales. Curr Biol, 28(23), 3850–3856 e9.

Waskom, M. L., Okazawa, G., & Kiani, R. (2019). Designing and interpreting psychophysical investigations of cognition. Neuron, 104(1), 100–112.

Witthoft, N., Sha, L., Winawer, J., & Kiani, R. (2018). Sensory and decision-making processes underlying perceptual adaptation. J Vis, 18(8), 10.

Witthoft, N., Poltoratski, S., Nguyen, M., Golarai, G., Liberman, A., LaRocque, K. F., Smith, M. E., & Grill-Spector, K. (2016). Reduced spatial integration in the ventral visual cortex underlies face recognition deficits in developmental prosopagnosia. bioRxiv, 051102.

Yamins, D. L., Hong, H., Cadieu, C. F., Solomon, E. A., Seibert, D., & DiCarlo, J. J. (2014). Performance-optimized hierarchical models predict neural responses in higher visual cortex. Proc Natl Acad Sci U S A, 111(23), 8619–24.

Yang, T., & Shadlen, M. N. (2007). Probabilistic reasoning by neurons. Nature, 447(7148), 1075–80.

Yin, R. K. (1969). Looking at upside-down faces. Journal of Experimental Psychology, 81(1), 141–145.

Young, A. W., Hellawell, D., & Hay, D. C. (1987). Configurational information in face perception. Perception, 16(6), 747–59.

Zhan, J., Ince, R. A. A., van Rijsbergen, N., & Schyns, P. G. (2019). Dynamic construction of reduced representations in the brain for perceptual decision behavior. Curr Biol, 29(2), 319–326 e4.

